# Associations of psychiatric disease and aging on *FKBP5* expression converge on cortical supragranular neurons

**DOI:** 10.1101/2021.01.27.428487

**Authors:** Natalie Matosin, Janine Arloth, Darina Czamara, Katrina Z. Edmond, Malosree Maitra, Anna Sophie Fröhlich, Silvia Martinelli, Dominic Kaul, Rachael Bartlett, Amber R. Curry, Nils C. Gassen, Kathrin Hafner, Nikola S Müller, Karolina Worf, Ghalia Rehawi, Corina Nagy, Thorhildur Halldorsdottir, Cristiana Cruceanu, Miriam Gagliardi, Nathalie Gerstner, Maik Ködel, Vanessa Murek, Michael J Ziller, Elizabeth Scarr, Ran Tao, Andrew E. Jaffe, Thomas Arzberger, Peter Falkai, Joel E. Kleinmann, Daniel R. Weinberger, Naguib Mechawar, Andrea Schmitt, Brian Dean, Gustavo Turecki, Thomas M. Hyde, Elisabeth B. Binder

**Affiliations:** Department of Translational Research in Psychiatry, Max-Planck Institute of Psychiatry, Munich, Germany; Illawarra Health and Medical Research Institute, Northfields Ave, Wollongong 2522, Australia; Molecular Horizons, School of Chemistry and Molecular Biosciences, Faculty of Science, Medicine and Health, University of Wollongong, Northfields Ave, Wollongong 2522, Australia; Institute of Computational Biology, Helmholtz Zentrum München, Neuherberg 85764, Germany; International Max Planck Research School for Translational Psychiatry, Munich, Germany; McGill Group for Suicide Studies, Douglas Mental Health University Institute, Montreal, Quebec, Canada; Department of Psychology, Reykjavik University, Reykjavik, Iceland; Neurohomeostasis Research Group, Institute of Psychiatry, University of Bonn, Clinical Centre, Bonn, Germany; Department of Psychiatry, McGill University, Montreal, Quebec, Canada; Department of Psychiatry, University of Münster, Münster, Germany; Melbourne Veterinary School, Faculty of Veterinary and Agricultural Sciences, The University of Melbourne, Parkville, Victoria 3010; The Lieber Institute for Brain Development, Johns Hopkins University Medical Campus, Baltimore, MD, USA; Department of Psychiatry and Psychotherapy, University Hospital, Ludwig-Maximilians University Munich, Nussbaumstrasse 7, 80336 Munich, Germany; Centre for Neuropathology and Prion Research, Ludwig-Maximilians University Munich, Nussbaumstrasse 7, 80336 Munich, Germany; Department of Psychiatry and Behavioral Sciences at Johns Hopkins University School of Medicine; Laboratory of Neuroscience (LIM27), Institute of Psychiatry, University of Sao Paulo, Rua Dr. Ovidio Pires de Campos 785, 05453-010 São Paulo, Brazil; Molecular Psychiatry Laboratory, Florey Institute for Neuroscience and Mental Health, Parkville, VIC, Australia; Department of Human Genetics, McGill University, Montreal, Quebec, Canada; Department of Psychiatry and Behavioral Sciences, Emory University School of Medicine, Atlanta, USA

**Keywords:** FKBP5, FKBP51, psychosis, depression, aging, stress, single-cell, postmortem brain

## Abstract

Identification and characterisation of novel targets for treatment is a priority in the field of psychiatry. *FKBP5* is a gene with decades of evidence suggesting its pathogenic role in a subset of psychiatric patients, with potential to be leveraged as a therapeutic target for these individuals. While it is widely reported that *FKBP5*/FKBP51 mRNA/protein (*FKBP5*/1) expression is impacted by psychiatric disease state, risk genotype and age, it is not known in which cell-types and sub-anatomical areas of the human brain this occurs. This knowledge is critical to propel *FKBP5*/1-targeted treatment development. Here, we performed an extensive, large-scale postmortem study (n=1024) of *FKBP5*/1 examining prefrontal cortex (BA9, BA11, BA24) derived from subjects that lived with schizophrenia, major depression or bipolar disorder. With an extensive battery of RNA (bulk RNA sequencing, single-nucleus RNA sequencing, microarray, qPCR, RNAscope) and protein (immunoblot, immunohistochemistry) analysis approaches, we thoroughly investigated the effects of disease-state, aging and genotype on cortical *FKBP5*/1 expression including in a cell-type specific manner. We identified consistently heightened *FKBP5*/1 levels in psychopathology and with age, but not genotype, with these effects strongest in schizophrenia. Using single-nucleus RNA sequencing (snRNAseq) and targeted histology, we established that these disease- and aging-effects on *FKBP5*/1 expression were most pronounced in excitatory supragranular neurons. We then found that this increase in *FKBP5* levels likely impacts on synaptic plasticity, as *FKBP5* gex levels strongly and inversely correlated with dendritic mushroom spine density and brain-derived neurotrophic factor (*BDNF*) levels in supragranular neurons. These findings pinpoint a novel cellular and molecular mechanism that has significant potential to open a new avenue of FKBP51 drug development to treat cognitive symptoms in psychiatric disorders.

## INTRODUCTION

Severe psychiatric disorders – including schizophrenia, major depression and bipolar disorder – present a huge burden for individuals, society and the healthcare system. While there are efficacious treatments, relapse, resistance and chronicity are common, and cognitive symptoms in particular are not adequately treated [17, 44]. A major issue is that the approved use of available drugs and treatments for specific psychiatric diagnoses, and their prescription to specific patients, are currently not based on biological knowledge of the pathological mechanisms that contribute to an individual’s disease presentation. Increased understanding of the molecular underpinnings of psychiatric disease, from the level of genes, to molecular and cell-type contributions, is critical to facilitate mechanism-based diagnoses, and the corresponding biological targets for treatment [50]. Despite decades of research, there are very few genes that have been translated from genetic association to a more mechanistic understanding of how this translates on the molecular and cellular levels and ultimately influences disease risk. This is an essential next step to both improve biological classification of psychopathology, and to drive treatment discovery.

FK506 Binding Protein 51 kDa (FKBP51), encoded by the *FKBP5* locus on chromosome 6p21.31, is a gene with strong evidence that it cross-diagnostically contributes to the pathogenesis of psychiatric disorders in a subset of patients. FKBP51 is an allosteric heat shock protein 90 kDa (HSP90) co-chaperone of the glucocorticoid receptor (GR), which is highly responsive to glucocorticoid (cortisol)-mediated stress. As such, *FKBP5*/1 is physiologically important for propagating and terminating the stress response [58]. While *FKBP5*/1 expression increases over the life course, even in healthy individuals [6, 10, 31, 51, 57], expression that exceeds this neurotypical rise during aging is likely to be harmful. Our previous studies indicate that amplified *FKBP5* expression is mechanistically caused by a combination of an *FKBP5* risk haplotype and lasting DNA methylation changes induced by early life adversity [23, 24]. In rodent models, heightened *FKBP5*/1 expression has been shown to cause psychiatric-like phenotypes, including impaired stress-coping behaviour, increased anxiety, and weakened extinction learning [58]. Evidence of this convergence of risk factors – disease-state, genotype, and age on *FKBP5*/1 expression – has been mainly distilled from clinical peripheral blood immune cells or rodent models. However how these factors coalesce in the human brain to raise risk to psychopathology or contribute to specific symptom domains is unknown, yet likely holds an important key to understanding *FKBP5*/1’s role in pathogenesis, as the brain is where many psychiatric symptoms manifest.

The extensive evidence of heightened *FKBP5*/1 expression as a risk factor for psychiatric disorders has spurred the development of small-molecule *FKBP5* antagonists as novel therapeutics [8, 31]. These agents have a promising profile of effects, with preclinical experiments in rodents showing that FKBP51 antagonists improve stress-coping behaviour and reduce anxiety when given systemically or directly into relevant brain regions such as the amygdala [8, 14]. To further fine-tune development of this drug-class, comprehensive characterisation of *FKBP5*/1 at the molecular level, with cell-type specificity, is urgently needed. Moreover, information on the status of *FKBP5*/1 directly in the human brain is required, given the brain is where psychotropic medications are most likely to exert their effects and that baseline differences in expression levels could alter the outcomes of a drug’s intended mechanism.

In the most comprehensive study to date, we examined a large collection (total n=1024) of human postmortem prefrontal cortex samples (Brodmann areas [BA] 9, 11 and 24) derived from individuals who lived with schizophrenia, major depression, or bipolar disorder compared to matched controls. These brain areas are highly implicated in psychiatric disorders, associated with hallmark symptoms of trans-diagnostic psychopathology such as executive functioning, emotion and working memory processes [1]. We used bulk and single-cell omics approaches and histological methods to explore the effects of disease-state, genotype and age on the cell-type- and cortical-layer-specific patterns of *FKBP5*/1 expression; importantly, we provide replication across 6 independent cohorts and robust validation of our results using multiple methods (e.g. seven different ways to assess *FKBP5*/1: bulk sequencing, single-nucleus sequencing, exon arrays, qPCR and RNAscope, immunoblot and immunofluorescent staining). Our results provide a new contribution to the field, indicating that effects of disease-state and age converge on supragranular neurons (superficial cortical layers LII-III), thus pinpointing a clear cellular target for further study and future drug development.

## MATERIALS AND METHODS

### Human postmortem brain samples

*FKBP5* gene expression (gex) and FKBP51 protein expression were examined in the human cortex from 1024 individuals across six postmortem brain cohorts (n=895 specimens from BA9, n=69 specimens from BA11, n=60 specimens from BA24) (Table 1). Informed consent was given by all donors or their next of kin; extensive details regarding collection, dissection and ethical approvals for each Cohort are included in the online resource (Online Resource, Supplementary Tables 1-6).

**Table 1.**
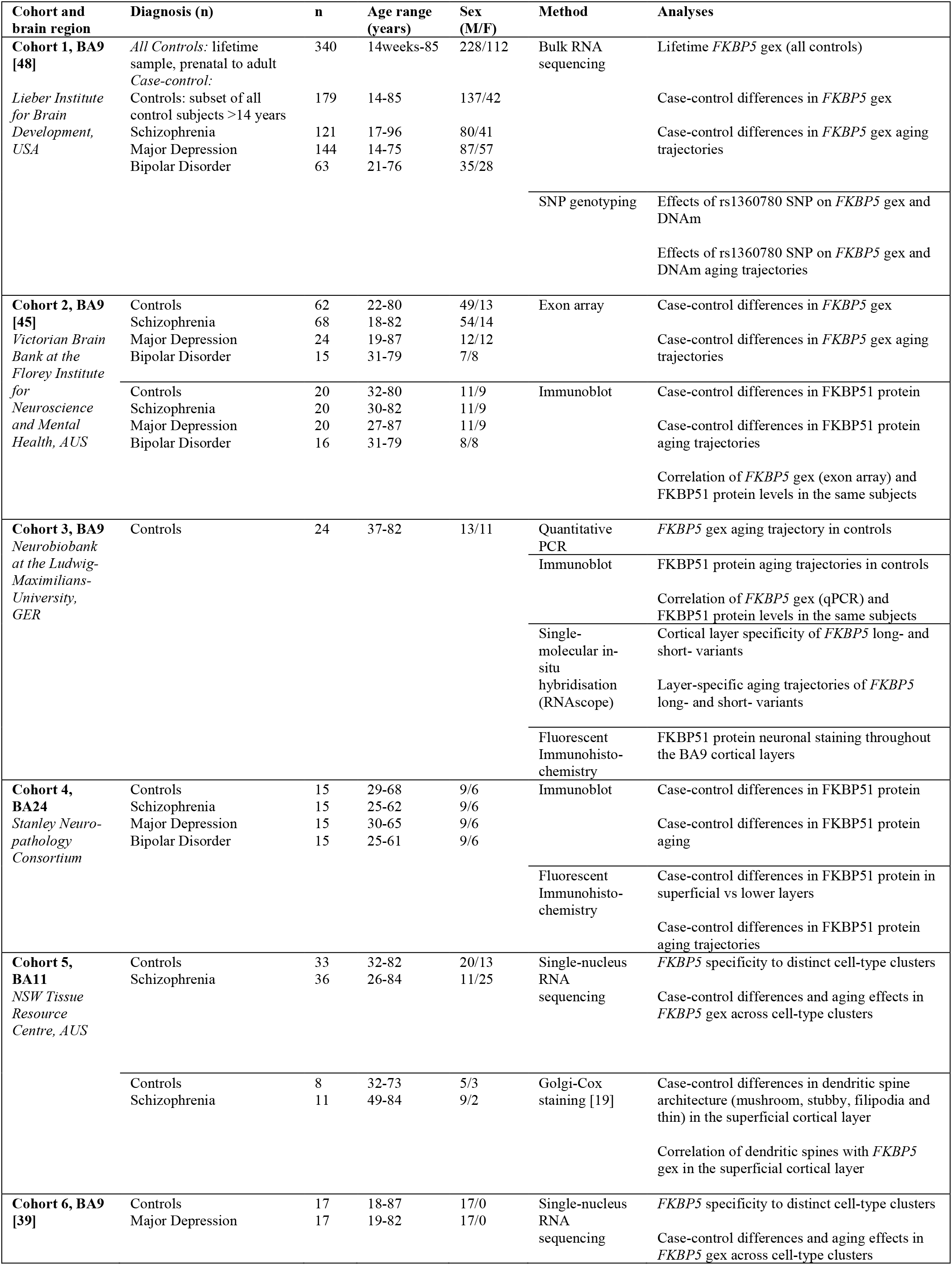
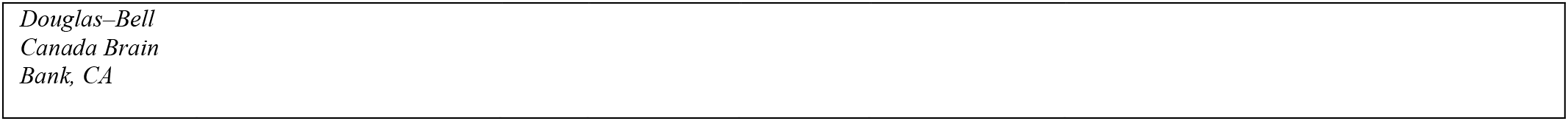
Summary of cohorts, methods, and analyses conducted.

### Bulk RNA sequencing (Cohort 1)

Bulk RNA sequencing methods have been previously described [48]. High-throughput sequencing was performed on the final cDNA library using the HiSeq 2000 (Illumina, San Diego, CA, USA), with the Illumina Real Time Analysis module used for image analysis and base-calling and the BCL Converter (CASAVA v1.8.2) to generate FASTQ files with sequencing pair-end 100 bp reads. Splice-read mapper TopHat (v2.0.4) was used to align reads to the human genome reference (UCSC hg19), with known transcripts provided by Ensembl Build GRCh37.67. Mapped reads covering the genomic region of *FKBP5* (chr6:35541362-35696397, GRCh37/hg19) were acquired. Reads covering each exon or unique exon-exon junction level were called using featureCounts (V1.5.0) [28]. Individual raw exon and junction reads were divided by the mapped total reads per subject and log normalised to account for skewedness (log2 fragments per kilobase million [FPKM]). We assessed reads covering a junction common to all *FKBP5* transcripts spanning from Exon 5 to Exon 6 (chr6:35565191-35586872).

### SNP genotyping (Cohort 1)

Genomic DNA from postmortem brain Cohort 1 was extracted from 100mg of pulverized cerebellum tissue with the phenol-chloroform method. SNP genotyping was performed with the HumaHap650Y_V3 or Human 1M-Duo_V3 BeadChips (Illumina) according to manufacturer’s instruction as previously described [48]. The rs1360780 *FKBP5* single nucleotide polymorphism (SNP) was extracted from the dataset. This SNP has been shown to affect *FKBP5* chromatin shape and transcription [24].

### Exon arrays (Cohort 2)

This method has been previously described [45]. Briefly, total RNA from Cohort 2 was isolated using TRIzol reagent (Life Technologies, Scoresby, VIC, Australia) and RNeasy mini kits (Qiagen, #74104, Chadstone Centre, VIC, Australia). RNA quality and quantity were assessed and samples with RINS of 7 or greater were deemed suitable for further analyses with the Affymetrix Human Exon 1.0 ST Array according to the manufacturer’s instructions (Affymetrix, Santa Clara, CA, USA). After hybridisation, chips were scanned and the fluorescent signals converted into a DAT file for quality control, and CEL and CHP files were generated. Exon array data was used as a replication for the RNAseq gex analyses in Cohort 1 and followed the same statistical methods. For *FKBP5* gex measures, we analysed signal from *FKBP5* exon 5 (ENSE00000747342.1) which sits adjacent to the *FKBP5* exon-exon junction used to assess total *FKBP5* gex in Cohort 1.

### Quantitative Real-Time PCR (Cohort 3)

mRNA levels of *FKBP5* transcripts in Cohort 3 were measured using qRT-PCR. 40mg of grey matter was dissected from each tissue block. RNA was extracted and RIN quantified as for RNA sequencing in Cohort 1. cDNA was synthesised from 500ng of RNA using the Maxima H Minus Reverse Transcriptase system (ThermoFisher Scientific). PCR conditions were set to 94°C for 5min, 40 cycles of 94°C for 30 s, 60°C for 30 s, 72°C for 30 s, and 72°C for 7min after the last cycle. Primer pairs were designed to amplify the unique junctions for total *FKBP5* gex (Online Resource, Supplementary Table 7), using customised TaqMan Gene Expression Assays (Applied Biosystems, Foster City, CA, USA) and the Lightcycler 480 (Roche, Basel, Switzerland). Gex levels of *FKBP5* was normalised to geometric means of the constitutively expressed genes β-actin (ACTB) and glyceraldehyde-3-phosphate dehydrogenase (GAPDH). Samples were measured in quadruplicate and averaged.

### Single-molecule fluorescent in situ hybridisation (RNAscope)(Cohort 3)

To quantify the gex and examine cortical localisation of *FKBP5* alternative transcripts, single-molecule fluorescent in situ hybridisation was performed on 16μm sections from Cohort 3 using the RNAscope Fluorescent V2 kit (Advanced Cell Diagnostics, Newark, CA, USA) according to the manufacturer’s instructions for the RNAscope® Part 1, Fresh Frozen Tissues protocol and the Multiplex Fluorescent Version 2 User Manual Part 2 (ACD). Two hybridisation probes were customised to target either *FKBP5* long transcripts (V1-3; 91000-2041bp) or *FKBP5* short transcript (V4; 1475-2968bp). Total *FKBP5* gex was calculated by summing these probes together. TSA fluorophores for cyanine 3 and cyanine 5 (Perkin Elmer) were used at a concentration of 1:1000. Slides were counter-stained with DAPI. Autofluorescence eliminator reagent (Merck Millipore-2160) was applied to reduce autofluorescence and slides were mounted with Aqua-Poly/Mount medium (Polysciences). Consistent regions of interest were identified based on gyri formation, with the anterior most point of each section imaged on a Leica SP8 confocal microscope. 11 z-stack images were taken at 0.5μm distance of the superficial (LII-III) and deep layers (LIV-VI) based on visual nuclei morphology, density and distance to the cortex edge or white matter. *FKBP5* long transcript probes were detected with cyanine 3 at 570nm, and the short transcript in cyanine 5 at 650nm. Optimal exposure time and image processing procedures were determined using positive controls (POL2RA for the long transcripts and PPIB for the short transcript), which were undetectable in sections hybridised with the negative control probe DapB. Fiji software [47] was used to merge the 11 z-stacks, adjust thresholds, and automatically quantify the number of nuclei and transcripts for each probe, with one dot representing one transcript. Number of dots were normalised to the number of nuclei, and the average for each subject was calculated.

### Immunoblot (Cohort 2, 3 and 4)

Relative protein densities of FKBP51 were determined by immunoblot in Cohort 2, 3 and 4. 10mg of grey matter from each block was homogenised and protein concentration determined using BCA assays. Samples were loaded in duplicate at a loading concentration of 20μg of total protein per lane and electrophoresed as previously described [30]. Membranes were then incubated overnight at 4°C with a validated primary antibody diluted with TBST containing 1% skim milk, followed by washing and secondary antibodies (Online Resource, Supplementary Table 8). Primary antibody specificities were validated in FKBP51 knock out (KO) cells, generated from the SH-SY5Y human neuroblastoma cell line using CRISPR-Cas9 (Online Resource, Supplementary Figure 1). Enhanced chemiluminescence was applied. The blots were visualised with the BioRad ChemiDoc XRS+ (Cohort 2) or the Amersham 6000 Gel Imager (GE Healthcare, Cohort 3), and quantified with Image Lab Software (BioRad). Membranes were re-probed and normalised to a loading control (Cohort 2: anti-β-actin polyclonal antibody Santa Cruz Biotechnology, Dallas, TX USA #sc-1616, 1:3000; Cohort 3: glyceraldehyde 3-phosphate dehydrogenase Abcam #ab9485, 1:2500). Densitometry values for each sample were normalised to the respective loading control, then to the respective pooled sample to account for gel-to-gel variability. Duplicates were averaged for each sample, and the mean of the two primary antibodies taken as the final values.

### Fluorescent Immunohistochemistry (Cohort 3 and 4)

To identify and semi-quantify cellular and subcellular localisation of FKBP51 protein, fluorescent immunohistochemistry was performed in Cohort 3 and 4. Fresh frozen sections (14-20μm) were post-fixed with 4% PFA and antigen retrieval performed using 10mM citric acid buffer with 0.5% Tween-20 for 10mins. Tissue was permeabilised with 0.3% Triton X-100 in phosphate buffered saline (PBS) for 5mins and blocked with 10% normal serum, 1% bovine serum albumin, with 0.3M glycine in PBS at room temperature for 1hr. Sections were incubated with primary antibody solution overnight at 4°C antibodies used are summarised in Online Resource, Supplementary Table 9. Following washing, appropriate secondary antibodies and DAPI were applied. Tissues were counterstained with autofluorescence eliminator reagent (Merck Millipore-2160) and cover-slipped with Aqua-Poly/Mount medium.

Wide-field microscopy images were acquired on the Leica THUNDER imager (Leica, Germany) and the Leica SP8 confocal microscope. 40x Z-stack images were acquired using a 1μm step-size at 512 x 512 pixels. Six stacks, collected at similar locations for each subject, with an average of 12 z-sections per stack were semi-randomly acquired within the deep and superficial grey matter region (three places each) of each section. Images were imported into ImageJ with the Fiji plugin [47] and merged using the *Z project* tool. The max projection images were imported into QuPath (Version 0.3.2), and *cell detection* and *show detection measurements* tools were used to quantify staining. The average intensity of FKBP51 staining per cell type was determined per image for both the FKBP51 detections and the background of the FKBP51 channel, and FKBP51 expression was normalised to the background to account for image-to-image variability. Analysis for Cohort 4 was performed using data acquired on the individual cell level with the data from each cell (e.g., FKBP51 staining intensity) taken as an individual data point and the subject that the cell came from was taken as a covariate [39]. With a dataset of this size (>6,000 cells), normality was assessed with histograms, boxplots and qqPlots using ggplot2 in R. Data was log transformed to achieve normal distribution. On average, data from one of the three acquired images from two subjects had to be excluded from each analysis due to being greater than ±2 standard deviations from the mean. Analysis was subsequently performed by using parametric tests.

### Cohort 5 and 6

#### Single-nuclei RNA sequencing

We performed snRNAseq in Cohort 5 with the 10x Genomics Chromium system (Single Cell 3’ Reagents kit v3.1) with 10,000 nuclei per sample as target recovery (~70,000 cells). Libraries were pooled equimolarly and were treated with Illumina Free Adapter Blocking Reagent before sequencing in two batches on the NovaSeq 6000 System (Illumina). Sequence reads were demultiplexed using the sample index, aligned to a pre-mRNA reference and UMI were counted after demultiplexing of nuclei barcodes using Cell Ranger v6.0.1. Reads were down-sampled per cell to the 75% quartile of reads per cell (14786 reads). Count matrices of all individuals were combined and further processed using Scanpy v1.7.1. Nuclei were filtered according to counts, minimum genes expressed and % of mitochondrial genes (counts <500, genes<300, Mito%≥15). Genes expressed in <500 nuclei were removed. Data were normalised and log-transformed using sctransform. Leiden clustering using highly variable genes was applied for clustering. A label transfer algorithm (scarches v0.4.0) was used for an initial cell type assignment. Thereby, cell type labels from the Allen Brain Atlas (Human M1 10x) were taken as a reference for our dataset. These initial assignments were refined by a manual curation based on marker gene expression [39, 49]. Cells were scaled to 10,000 reads (raw counts), and *FKBP5* expression values per single cell were extracted and averaged per subject to obtain a pseudo cell per subject per identified cell type. Excitatory neuron sub-clusters Ex L4-6(2), Ex L5-6(2), and Ex 20 were removed due a high number of zero-expressors which skewed the data, and an inability to determine if these were biological or technical dropouts. We also analysed Cohort 6 10x Genomics Chromium data (v2 chemistry) previously described [39]. Ex 1, Ex 5 and Ex 9 were also removed due a high number of zero-expressors. Data were processed with Cellranger and Seurat to identify cell-type clusters. The FindAllMarkers function was used with the bimodal test and logfc.threshold of log(2), with other parameters set to default. *FKBP5* expression per single cell was extracted. All cells were scaled to 10,000 reads and then averaged per subject to create a pseudo cell per subject per classified cell-type in the dataset.

#### Golgi-Cox staining

Golgi-Cox staining was performed in a subset of Cohort 5 (BA11) as previously described [19]. Fresh frozen tissue blocks (~0.5cm) were stained using the FD Rapid GolgistainTM Kit following the manufacturer’s instructions (FD Neurotechnologies, Columbia, MD, USA). Microscopy was completed using brightfield imaging (DMi8; Leica Microsystems, Wetzlar, Germany). Cortical layers I-VI (LI-VI) were defined according to morphological features [7] and dendritic spines on supragranular (superficial) and infragranular (deep layer) pyramidal neurons were measured using an optical fractionator method to randomly select coordinates within each layer. This was repeated until 3–6 pyramidal neurons were imaged in the supragranular and infragranular layers. A total of 12,144 dendritic spines were quantified in LII/III of BA11 in Cohort 5 and analysed for correlation with superficial layer *FKBP5* gex snRNAseq data.

### Statistics

Analyses were performed in R v3.3.1 (https://www.r-project.org). A summary of the statistics implemented, has been included in the Online Resource, Supplementary Table 10. All reported *P* values from *lm* analyses were calculated from *t* statistics computed from the log fold change and its standard error from each multiple regression model. To account for multiple-testing, False Discovery Rate (FDR) correction was computed for all P-values obtained comparing different diagnoses (schizophrenia/depression/bipolar vs control) within each statistical test, using an *a priori* standard FDR cut-off value of 0.05 [3]. To account for the increased chance of type 1 error due to many datapoints [12], we considered only significant FDR-corrected P values with effect sizes that are likely to have biological significance (correlation R>0.25 and mean expression differences >10%).

## RESULTS

### *FKBP5* mRNA and FKBP51 protein levels are higher in schizophrenia subjects

To comprehensively understand how expression patterns of *FKBP5* gex and FKBP51 protein are altered in severe psychopathology, we explored the largest sample to date to assess *FKBP5* cortical expression differences in cases compared to controls (see Table 1 for explanation of postmortem cohorts and sample sizes). In Cohort 1 we observed a striking +28.0% mean expression difference in *FKBP5* gex in all cases vs controls measured by RNAseq (Cohort 1, n=507: t=3.299, *P*=0.001043,). Post-hoc analysis revealed that this was driven by +39.8% higher *FKBP5* gex in subjects with schizophrenia (t=3.226, *P*=0.001403, FDR=0.004209) and depression (+23.86%; t=2.478, *P*=0.01377, FDR=0.020655) (Figure 1a). We replicated this finding in an independent sample (Cohort 2, n=169, see Table 1) using a different method (exon arrays), with increased in *FKBP5* gex in cases vs controls overall (t=3.325, *P*=0.001117) again driven by the schizophrenia subjects (t=3.348, *P*=0.00112, FDR=0.00336) (Figure 1b). At the protein level, we found FKBP51 protein was also increased specifically in schizophrenia vs controls (Cohort 2: t=2.580, *P*=0.0149, FDR=0.04470, +17.3%, Figure 1c; Cohort 4: t=2.685, *P*=0.00972, FDR=0.02916, +18.3%, Figure 1d). This was consistent with a strong, positive correlation between *FKBP5* gex and FKBP51 protein levels across all subjects in Cohort 2 (R=0.507, *P*=4.57E-05, Figure 1e). Altogether these results support that *FKBP5* gex and protein levels are increased in psychiatric cases vs controls, with the most pronounced effects seen in schizophrenia subjects.

**Figure 1.**
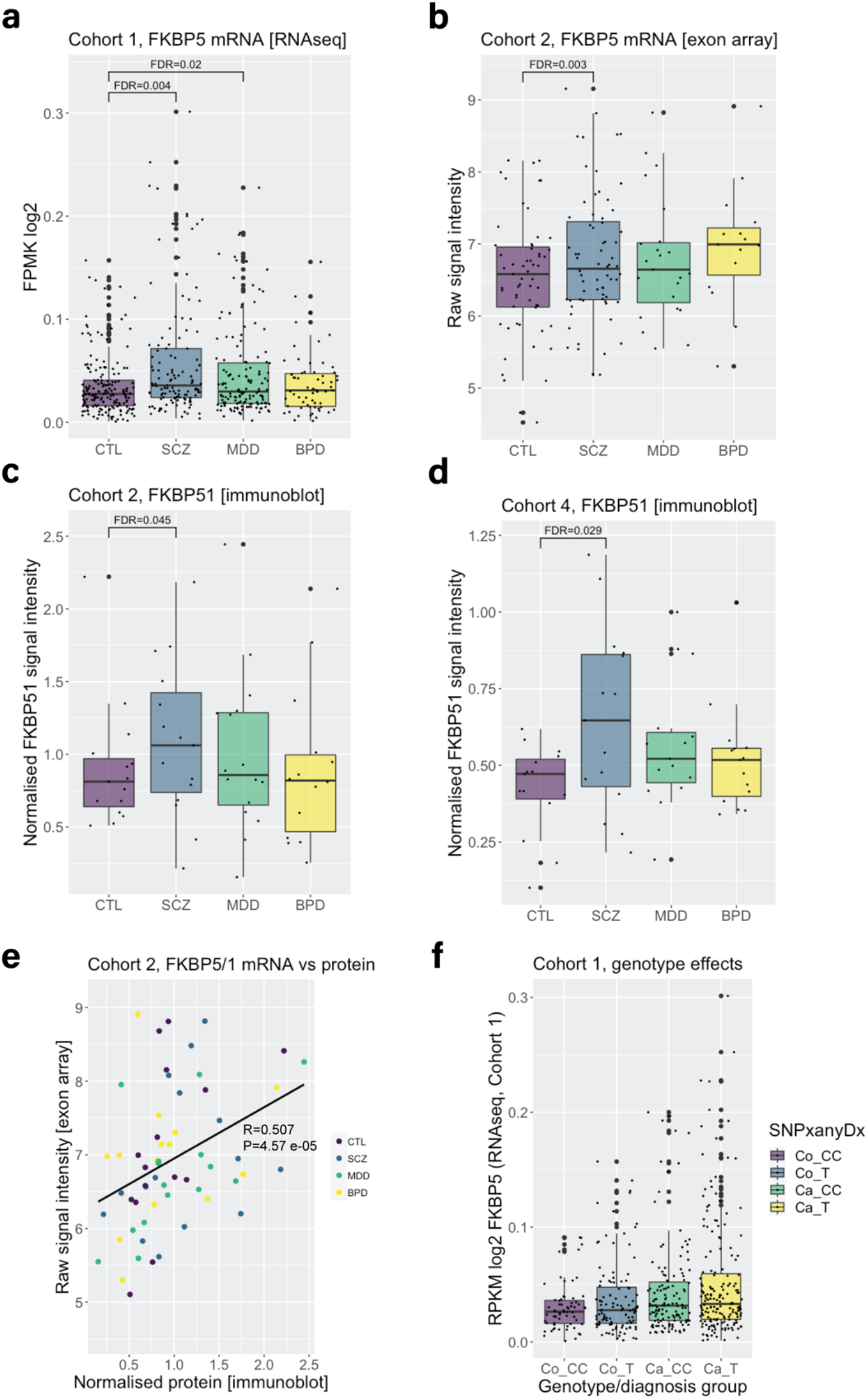
Differences in *FKBP5*/FKBP51 expression in schizophrenia, major depression and bipolar disorder vs controls. **(a)** In Cohort 1, linear regression modelling revealed *FKBP5* expression was significantly increased in subjects with schizophrenia and major depression compared to controls. Gene expression in Cohort 1 was measured with bulk RNA sequencing assessing reads covering an exon-exon junction between exon 5 and exon 6, common to all *FKBP5* alternative transcripts (chr6:35565191-35586872). **(b)** In Cohort 2, linear regression modelling revealed that *FKBP5* expression was significantly increased in subjects with schizophrenia compared to controls. Gene expression in Cohort 2 was measured with exon arrays assessing signal intensity from *FKBP5* exon 5 (ENSE00000747342.1) which sits adjacent to the *FKBP5* exon-exon junction used to assess *FKBP5* with RNAseq in Cohort 1. **(c)** Linear regression modelling revealed FKBP51 protein expression in Cohort 2 was increased in schizophrenia subjects compared to controls. FKBP51 protein expression was measured with immunoblot, with FKBP51 signal intensity normalised to a loading control and pool to account for sample-to-sample and gel-to-gel variability. **(d)** Linear regression modelling revealed FKBP51 protein expression in Cohort 4 was increased in schizophrenia subjects compared to controls. FKBP51 protein expression was measured with immunoblot, with FKBP51 signal intensity normalised to a loading control and pool to account for sample-to-sample and gel-to-gel variability. **(e)***FKBP5* mRNA and FKBP51 protein expression in Cohort 2 were significantly and positively correlated (all subjects analysed together irrespective of diagnosis, Spearman’s correlation). (**f**) In Cohort 1 (RNAseq) we observed a main effect of case-status on *FKBP5* mRNA expression (B=0.01144, t=2.989, *P*=0.0.00295), but there was no additive effect of genotype (B=0.00144, t=−0.592, *P*=0.55399). *Abbreviations*: Co, control; Ca, Case; CC, CC homozygotes and T, T carriers. “SNPanyDx” indicates that this was across all cases combined. *Abbreviations*: BPD, bipolar disorder; CTL, control; gex, gene expression; FDR: false discovery rate corrected P values (to account for multiple comparisons); MDD, major depressive disorder; SCZ, schizophrenia; Cohorts detailed in Table 1.

### No effect of *FKBP5* risk genotype on heightened *FKBP5* mRNA levels

Previous studies have reported that *FKBP5* gex is differentially induced by glucocorticoids based on *FKBP5* rs1360780 genotype, with minor allele (T) carriers exhibiting the highest levels of *FKBP5* gex after such stimulation [21, 36]. We tested for additive effects of this genotype and case-status on *FKBP5* gex in Cohort 1 in all cases (combined to improve power) vs controls (T carrier/CC, control n=108/64, schizophrenia n=69/45, major depression n=78/62, bipolar disorder n=32/28; Online Resource, Supplementary Table 11). We observed a main effect of case-status on *FKBP5* gex (B=0.01144, t=2.989, *P*=0.0.00295), but there was no additive effect of genotype (B=0.00144, t=−0.592, *P*=0.55399; Figure 1f).

### *FKBP5* mRNA and bulk FKBP51 protein levels increase with age over the human lifespan

Given age is an important modifier of disease presentation and treatment outcome [34], and reported to impact on *FKBP5*/1 expression [6, 31, 51], we set out to investigate aging effects on *FKBP5***/** 1 in our cohorts. We firstly characterised the expression pattern of *FKBP5* throughout life by analysing a large sample of neurotypical controls from 14 weeks gestational age to 85 years (Cohort 1; n=340; Figure 2a). We found that *FKBP5* gex naturally inflected in childhood and adolescence and increased consistently from 14 to 88 years of age (R=0.424, *P*=2.2E-16, Figure 2a). The strong, positive correlation of gex with age in control adults was also seen in Cohort 2 (Online Resource, Supplementary Figure 2a; R=0.61343, *P*=1.153E-07) and validated with qPCR using two probes targeting total *FKBP5* gex in Cohort 3 (Probe 1, R=0.460, *P*=0.024; Probe 2, R=0.449, *P*=0.028; Online Resource, Supplementary Figure 2b-c). FKBP51 total protein expression measured with western blot (bulk tissue) was also positively correlated with age in control adults (Cohort 2: R=0.593, *P*=0.022; Cohort 3: R=0.500, *P*=0.014; Figure 2b). In Cohort 4, FKBP51 and age were not significantly correlated (R=0.483, P=0.639; Online Resource, Supplementary Figure 2d). This is likely because Cohort 4 is a younger cohort with only n=4 subjects over 55 years of age.

**Figure 2.**
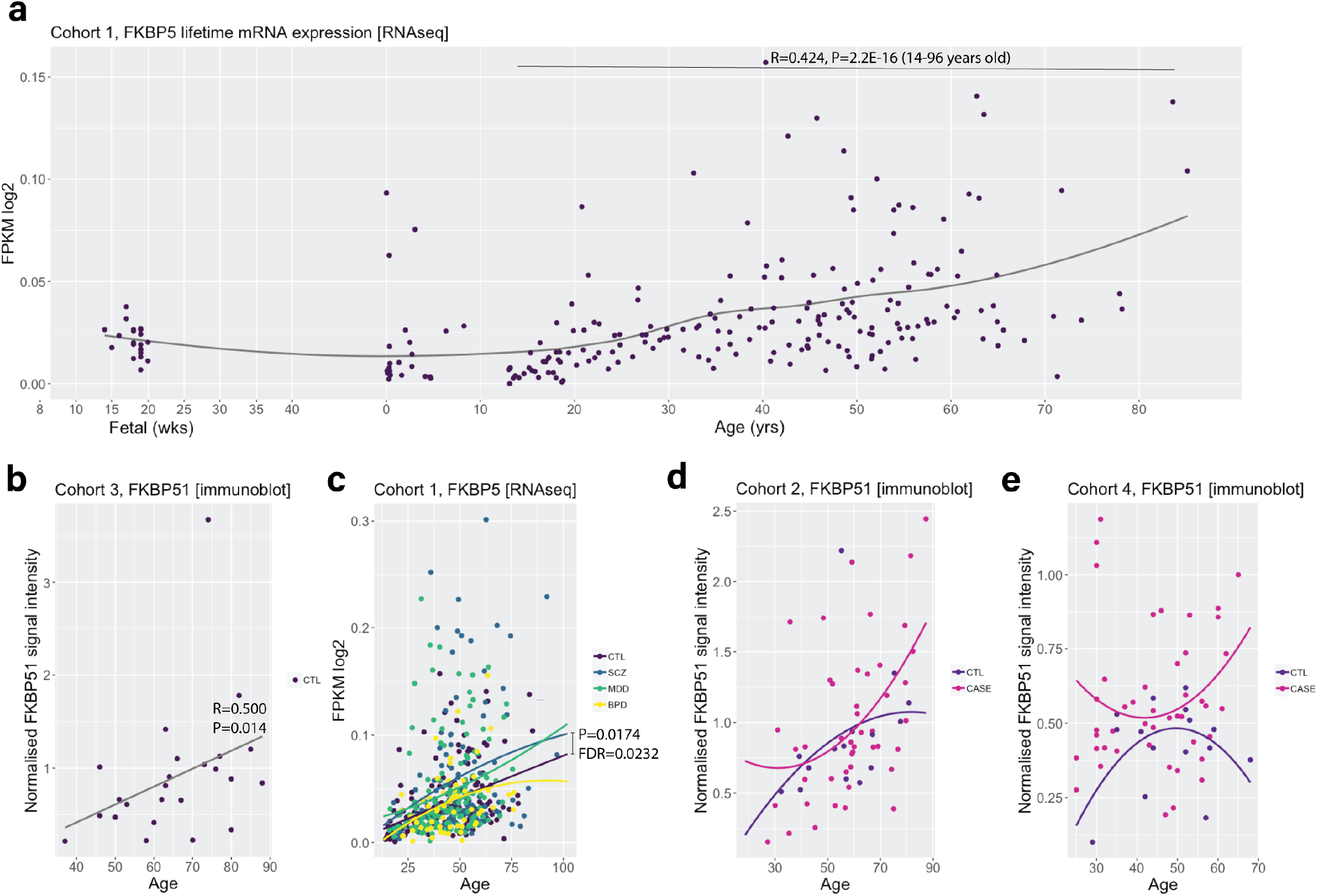
Effects of age on FKBP5/1 expression levels over the lifetime and in major psychiatric illnesses. **(a)** Trajectory of *FKBP5* gene expression in control subjects over the lifespan, from foetal time points to 88 years of age (Cohort 1, loess fit curve). *FKBP5* gene expression increased over the life course in healthy subjects, and was positively correlated with age between 14-96 years of age (calculated with Spearman’s correlations). **(b)** FKBP51 protein expression was positively correlated with age in Cohort 3 (calculated with Spearman’s correlations). **(c)***FKBP5* aging trajectory was significantly heightened in schizophrenia subjects vs controls (Cohort 1, comparison of nonparametric curves achieved using sm.ancova statistics). **(d-e)** FKBP51 protein aging trajectories were not significant in cases vs controls in Cohorts 2 or 4. The plots show a comparison of nonparametric curves achieved using sm.ancova statistics. *Abbreviations*: BPD, bipolar disorder; CTL, control; MDD, major depressive disorder; SCZ, schizophrenia; Cohorts detailed in Table 1.

### *FKBP5*/FKBP51 aging effects are further increased in psychiatric disorders

We then examined the effects of age on *FKBP5* gex in cases compared to controls (subjects >14 years), to determine if having a psychiatric disorder alters the neurotypical increase in *FKBP5* gex with age. We initially determined if there was a significant increase in the aging trajectory by comparing the correlation slopes with *sm.ancova*. In Cohort 1, *FKBP5* gex was more positively correlated with age in schizophrenia subjects vs controls and the aging trajectory was significantly heightened in the schizophrenia subjects compared to controls (*P*=0.0174, FDR=0.0232; Figure 2c). To quantify the effect size of the expression difference at older ages, we calculated the mean expression difference in schizophrenia vs control subjects over 50, which was +29.5% higher compared to neurotypical controls over 50 (t=2.060, *P*=0.02593, FDR=0.050, Figure 2c). In Cohort 2, no significant difference in the schizophrenia vs control *FKBP5* mRNA aging trajectory was found (*P*=0.056; Online Resource, Supplementary Figure 2a).

At the protein level in Cohorts 2 and 4, there was not enough statistical power to accurately compare the aging trajectories in each diagnosis vs neurotypical controls with *sm.anova* due to too few cases in each diagnosis at older ages. However, given the strong correlation of *FKBP5* gex and FKBP51 protein (Figure 1e), it is likely that the pronounced increase in mRNA with age in schizophrenia also occurs at the protein level. Indeed, the highest FKBP51 protein expression was observed in older cases in both Cohort 2 (Figure 2d; subjects over 50: case protein 0.971±0.464 vs control protein 0.885±0.388, +9%) and Cohort 4 (Figure 2e; subjects over 50: case protein 0.612±0.201 vs control protein 0.442±0.132, +27%). These results collectively support that older cases have higher *FKBP5* mRNA and likely FKBP51 protein levels compared to neurotypical controls.

### Localisation of *FKBP5*/FKBP51 expression in the prefrontal cortex

To gain deeper insight into the cell-type specificity of heightened *FKBP5*/1, we assessed *FKBP5*/1 cell-type distribution and the cortical-layer specificity. To date, there is little known about the expression patterns of the *FKBP5*/1 in the human cortex including its cell-type and cortical-layer distribution. We firstly used single-nucleus RNA sequencing (snRNAseq; Cohort 5 and 6) to examine the levels of *FKBP5* gex according to cell-type (Figure 3a-d). Among the 20 cell-type clusters in Cohort 5 (BA11, n=69; Figure 3a) and 26 delineated cell-type clusters in Cohort 6 (BA9, n=34; Figure 3b), *FKBP5* gex was most highly expressed in excitatory neurons, microglia, and astrocytes (Figure 3c-d). We also compared our results with other publicly available human prefrontal cortex snRNAseq datasets [13, 26] (Figure 3e). In BA9 (n=3) [13], expression was highest in excitatory neurons and microglia. In the adjacent BA6/BA10 areas (n=6) [26], the highest levels of *FKBP5* were observed in excitatory and inhibitory neurons. High *FKBP5* gex in excitatory neurons was thus a consistent finding across the four cortical snRNAseq datasets.

**Figure 3.**
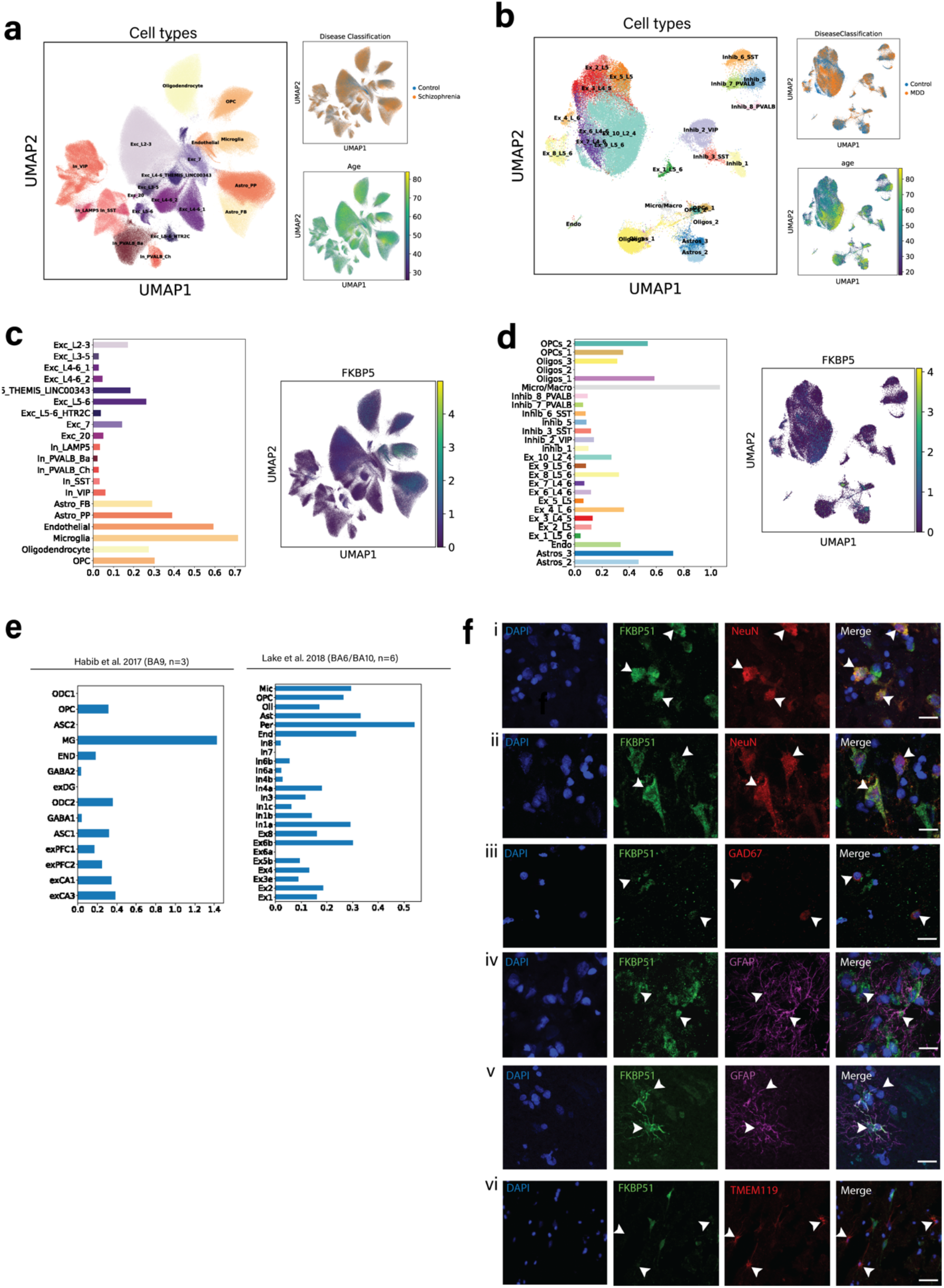
Cell-type distribution of FKBP5/FKBP51 expression in the prefrontal cortex (BA9, BA10, BA6, BA11). In **(a)** Cohort 5, and **(b)** Cohort 6, dimensionality reduction uniform manifold approximation and projection (UMAP) plots depicting total cells across all individuals. The colours indicate the delineated sub-cell type clusters (left of each panel), the distribution of diagnosis across the clusters (top right), and the distribution of age across clusters (bottom right). In **(c)** Cohort 5 and (**d**) Cohort 6, bar plots show the average *FKBP5* gene expression per cell-type cluster, as well as *FKBP5* expression across cell clusters. (**e**) Bar plots showing the average FKBP5 gene expression per cell type cluster in previously published datasets [13, 26] **(f)** Co-localisation of FKBP51 protein expression (green) with nuclear marker DAPI (blue) and: (i) NeuN+ neurons (red) in the supragranular layer; (ii) NeuN+ neurons in the infragranular layer; (iii) GAD67+ neurons; (iv-v) GFAP+ astrocytes (top showing no colocalization and bottom showing strong colocalization); (vi) TMEM119+ microglia. Scale bar for all images = 20μm.

Fluorescent immunohistochemistry (Cohort 3) confirmed high FKBP51 protein expression in the soma and nucleus of both NeuN+ (Figure 3f, i-ii supragranular/infragranular respectively) and GAD67+ (Figure 3f, iii) cortical neurons. FKBP51 staining intensity was lower and more inconsistent on glia. For example, GFAP+ astrocytes had highly variable levels of FKBP51 expression, with some cells showing little visually detectable FKBP51 expression (Figure 3f, iv), and others showing high expression throughout the entire astrocyte (Figure 3f, v). We also found FKBP51 was localised on some but not all TMEM119+ microglia, albeit with lower expression than in neurons (Figure 3f, vi).

To localise these observations to cortical layers, entire tilescan images of FKBP51 staining were evaluated (Cohort 3 and neurotypical controls of Cohort 4, Figure 4a-b). In BA9 (Cohort 3), FKBP51+ cells (all cell-types) were observed both in the superficial layers (LII-III) the deeper layers (55% in superficial and 71% in deep, Figure 4c, i). Additionally, 80% of superficial layer NeuN+ neurons and 84% of deep layer were FKBP51+ (Figure 4c, ii). High levels of FKBP51 expression but with somewhat different distribution was also observed in BA24 (Cohort 4), with 91% of cells (all types) in the superficial layer being FKBP51+ and 64% in the deeper layers (Figure 4c, iii). Additionally, 65% of superficial and 72% of deep layer NeuN+ neurons were FKBP51+ (Figure 4c, iv). It was not possible to accurately quantify the proportions of FKBP51+ glia due to the inconsistent staining on cell bodies and processes. Overall, snRNAseq and immunohistochemistry both support that the most consistent and intense *FKBP5*/1 gex/staining was localised on excitatory neurons with slightly varied expression across cortical layers.

**Figure 4.**
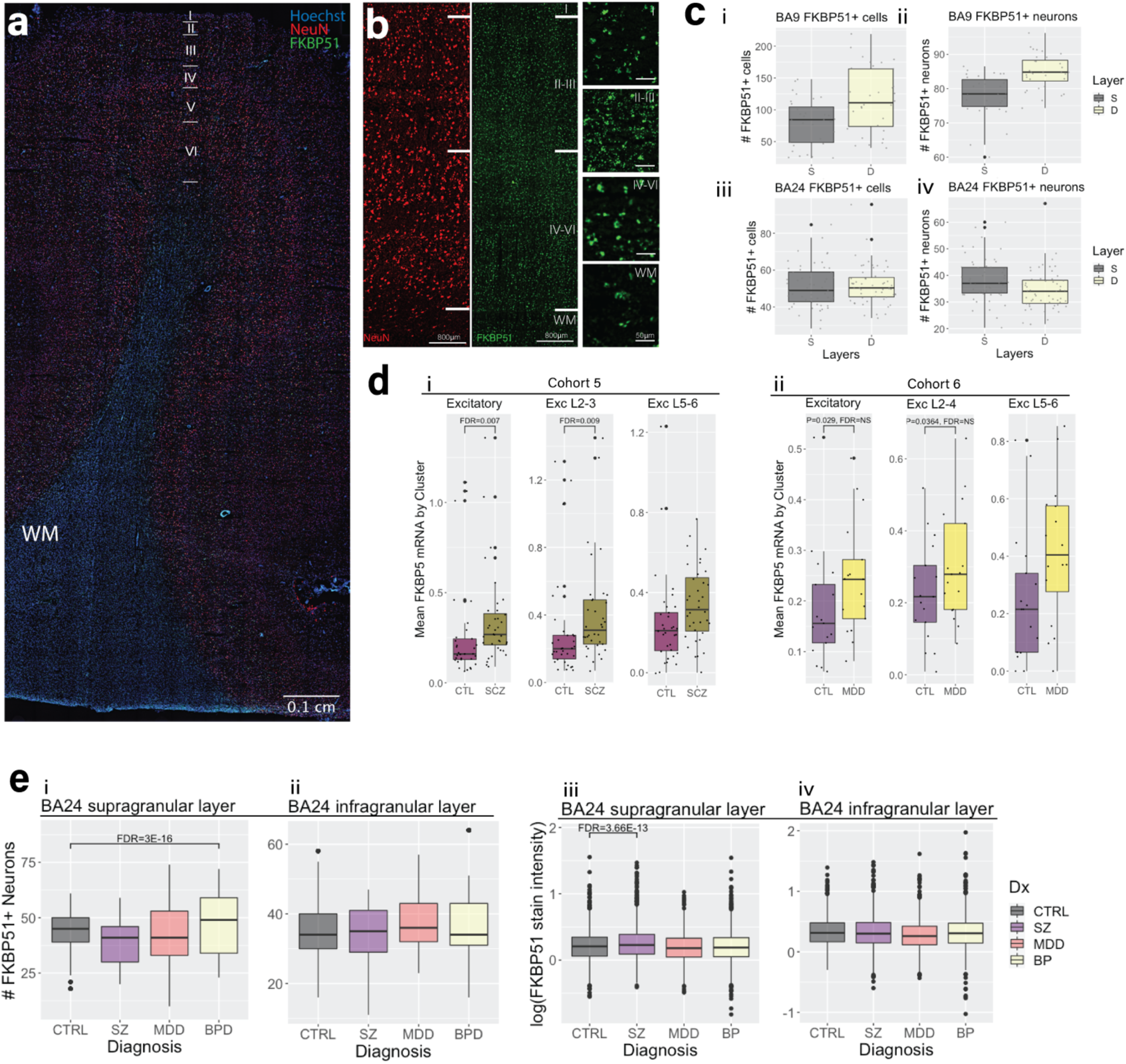
Case-control differences in *FKBP5*/1 levels in supragranular vs infragranular cortical layer excitatory neurons. **(a)** Representative tilescan image showing FKBP51 staining in a section of BA9 cortex. **(b)** Same image at higher magnification showing the pattern of NeuN neuronal staining (red) and FKBP51 staining (green) across the cortical layers; Layers II-III denote the supragranular cortex, layers IV-VI denote the infragranular cortex and WM denotes the white matter. **(c)** Box plots showing the number of FKBP5+ cells (left) and neurons (right) in BA9 (i-ii) and in BA24 (iii-iv). S denotes the superficial or supragranular layer, and D denotes the deep or infragranular layer. **(d)** snRNAseq data from Cohort 5 (i) and Cohort 6 (ii) showing FKBP5 gene expression levels in the excitatory neuron group (total, left), excitatory neurons from cortical layer 2-4 (Exc 2-4, middle) and layer 5-6 (Exc 5-6, right) in control (CTL) vs schizophrenia (SCZ) and major depressive disorder (MDD). FKBP5 expression was significantly higher in schizophrenia, but not depression, vs controls after correcting for multiple comparisons. Data was analysed using linear regression modelling. **(e)** Boxplots of immunohistochemistry data demonstrating the number of FKBP51+ neurons in (i) the supragranular layer and (ii) the infragranular layer in each psychiatric disorder, as well as the intensity in the supragranular (iii) and infragranular layers in each psychiatric disorder. All differences illustrated with boxplots were calculated with linear regression modelling. *Abbreviations*: CTRL, controls, SZ, schizophrenia, MDD, major depressive disorder, BP, bipolar disorder.

### Cell-type specificity of heightened FKBP5/1 expression in psychopathology

We next assessed case-control differences in *FKBP5* gex within our single-cell-type clusters. In Cohort 5 (BA11, schizophrenia vs control n=69), *FKBP5* gex was higher in schizophrenia vs controls in all cell type groups except inhibitory neurons (output of all comparisons in Online Resource, Supplementary Table 12). Given our finding of strong co-localisation of FKBP5/1 specifically on NeuN+ pyramidal neurons, we further examined case-control differences in *FKBP5* gex in the excitatory neuron sub-clusters. *FKBP5* gex was substantially higher in the excitatory neuron group in schizophrenia (+32%, t=2.969, *P*=0.00426, FDR=0.00746, Figure 4d, i). Subsequent analysis of *FKBP5* gex in the excitatory neuron sub-clusters showed that this increase was specific to the Ex 2 cluster representing LII-III excitatory neurons, with a striking +37% higher *FKBP5* gex in these neurons in schizophrenia vs controls (t=3.256, *P*=0.00186, FDR=0.0093, Figure 4d, i). In Cohort 6 (BA9, depression vs control, n=34), *FKBP5* gex expression was +24.8% higher in depression in the excitatory neuron group, although this did not survive correction for multiple cluster comparisons (t=−2.327, *P*=0.0267, FDR=0.187; Figure 4d, ii; Online Resource, Supplementary Table 13 and Supplementary Figure 3). This increase was again exclusive to superficial LII-IV excitatory neurons (Ex 10 cluster; +25%), although the comparison also did not survive multiple correction (t=−2.190, *P*=0.0364, FDR=0.255; Figure 4d, ii). There were no other significant depression/control differences between *FKBP5* gex in any other cell-type groups, suggesting the heightened *FKBP5* gex observed in schizophrenia and possibly to a lesser extent in depression was specific to supragranular excitatory neurons.

To validate the excitatory neuron finding at the protein level, we examined Cohort 4 with semi-quantitative immunohistochemistry examining supragranular vs infragranular FKBP51+ NeuN+ neurons in cases vs controls. Count and staining intensity data was collected for >12,000 NeuN+ neurons across all cortical layers. We first examined count data. In the supragranular layer, there was no difference in the number of FKBP5+ NeuN+ neurons in schizophrenia or depression, but there was an increase in bipolar disorder vs controls (+10% mean difference; t=10.683, *P*=2.00E-16, FDR=3.00E-16, Figure 4e, i). There was no difference in the number of FKBP51+ NeuN+ neurons in the infragranular layer (Figure 4e, ii). We next examined intensity data. The intensity of FKBP51 staining on supragranular NeuN+ neurons was +11.4% higher in schizophrenia subjects vs controls (t=−7.408, *P*<1.22E-13, FDR<3.7E-13; Figure 4e, iii); this increase was not seen in the infragranular layer (Figure 4e, iv). In depression, the intensity of FKBP51 expression was −13.1% lower throughout both the superficial and deeper layers (superficial: t=−9.273, *P*<0.001, FDR<0.001, vs. deep: t=− 4.907, *P*<0.001, FDR<0.001, −18.30%; Figure 4e, iii-iv). There was no difference seen in FKBP51 staining intensity in bipolar disorder. These findings consistently suggest that the increase in *FKBP5*/1 is specific to cortical excitatory neurons of the supragranular. layer, with the most pronounced differences observed in schizophrenia subjects.

### Age influences *FKBP5* and FKBP51 expression in superficial-layer excitatory neurons

We next checked for aging effects of *FKBP5* gex cell-type specifically, by correlating *FKBP5* gex and age in all major cell-type clusters (cases and controls combined). In Cohort 5, age was correlated with *FKBP5* gex in all the major cell-type clusters (Online Resource, Supplementary Table 14). However, the strongest correlation was observed in the excitatory neuron cluster (R=0.554, *P*= 8.02e-07, FDR=5.6E-06), and assessment of the excitatory neuron subclusters revealed that the strongest effect was in the superficial layer II-III excitatory neuron cluster (R=0.555, *P*=7.3e-07, FDR=3.65E-06, Figure 5a, i). We similarly observed a strong, positive association of *FKBP5* gex with aging in the excitatory neuron cluster in Cohort 6 (R=0.664, *P*=1.866e-05, FDR=5.6E-06), and again, assessment of the excitatory neuron subclusters revealed that the aging effect was strongest in, and specific to, the superficial layer 2-4 excitatory neurons (R=0.700, *P*=4.109e-06, FDR=2.88E-05, Figure 5a, ii, Online Resource, Supplementary Table 15). RNAscope was used to validate these findings histologically by accurately quantifying the aging-related changes of *FKBP5* transcripts in the superficial vs the deep layers of the BA9 cortex in Cohort 3. Our results also showed a positive correlation of *FKBP5* transcript dots and age in the superficial layers of BA9 (R=0.427, *P*=0.018, FDR=0.038), but not the deep layers (R=−0.037, *P*/FDR=0.864; Figure 5b). In Cohort 4, we were unable to confirm aging effects with immunohistochemistry as there were too few subjects at older ages (see Online Resource, Supplementary Table 16). However, in Cohort 3, FKBP51 staining intensity was 2.9-fold higher at 88 years compared to 37 years (Figure 5c). Altogether these findings consistently show the strong effect of age on *FKBP5*/1 gene and protein expression is pronounced in superficial layer excitatory neurons.

**Figure 5.**
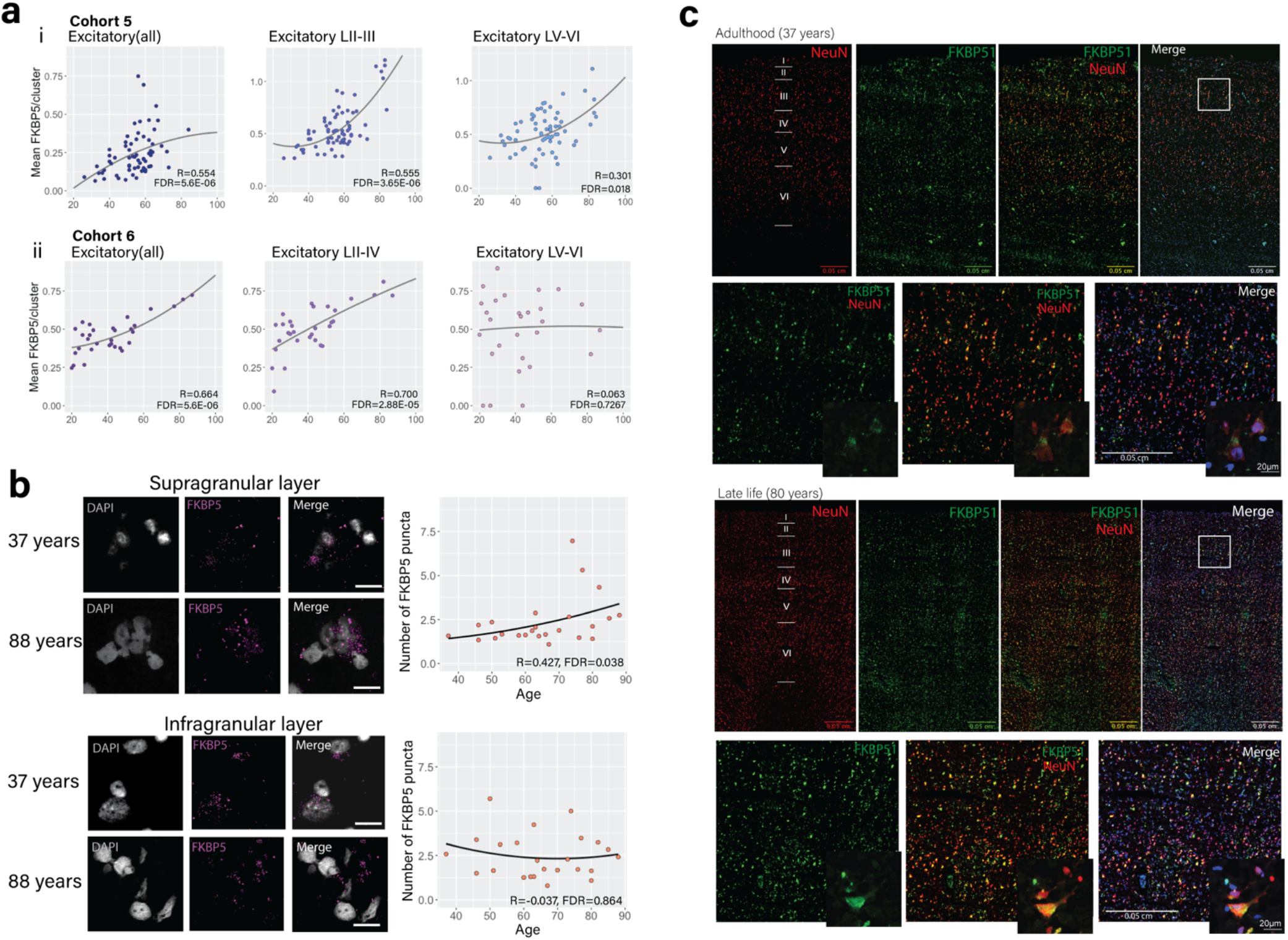
Layer-specific patterns of FKBP51 expression across the prefrontal cortex (BA9). **(a)** snRNAseq data showing *FKBP5* gene expression vs age in excitatory neurons (all, left), supragranular (LII-III, middle) and infragranular (LIV-VI, left) neuron clusters in (i) Cohort 5 (BA11), and (ii) Cohort 6 (BA9) using Spearman correlations. **(b)** RNAscope representative images (left) and results of quantification and analysis with Spearman correlations (right) showing that there was a positive association between *FKBP5* gene expression (purple puncta, scale bar=20μm) and age in the supragranular layer (top) but not the infragranular layer (bottom). **(c)** FKBP51 immunostaining across the BA9 cortex using widefield microscopy in adulthood (top panel, 37 years old), and later in life (bottom panel, 80 years old). Red represents staining with the neuronal markers NeuN, green represents staining of the FKBP51, and blue represents nuclear staining (Hoechst). The bottom images from each panel are at higher magnification taken from the white rectangle in the top right image. These images demonstrate FKBP51 staining is both high in neurons, and also is more intense (representing higher expression) at older ages.

### Impacts of elevated FKBP51 expression on dendritic spine architecture

Elevated *FKBP5* in the brain has been reported to suppress of glucocorticoid-receptor-mediated synaptic plasticity via impacts on dendritic architecture [4, 29, 53]. We therefore determined if the elevated *FKBP5* gex levels in superficial neurons in schizophrenia (Cohort 5, snRNAseq gex) correlated with changes in dendritic spine architecture in the same subjects with Golgi-Cox staining and quantification of mushroom, stubby, filopodia and thin dendritic spine subtypes (Figure 6a-c). While there was a negative correlation of *FKBP5* gex levels with dendritic spines (all types), this did not reach statistical significance (R=−0.3052136, *P*=0.2038). However further investigation of individual dendritic spine classes revealed *FKBP5* gex levels were strongly inversely correlated with mushroom spine density (R=−0.654, *P*=0.002378, FDR=0.009512; Figure 6d). These data suggest that the increased levels of *FKBP5* gex in superficial layer cortical neurons we consistently observed throughout our study may be related to a decrease in the number of mushroom dendritic spines on supragranular neurons.

**Figure 6.**
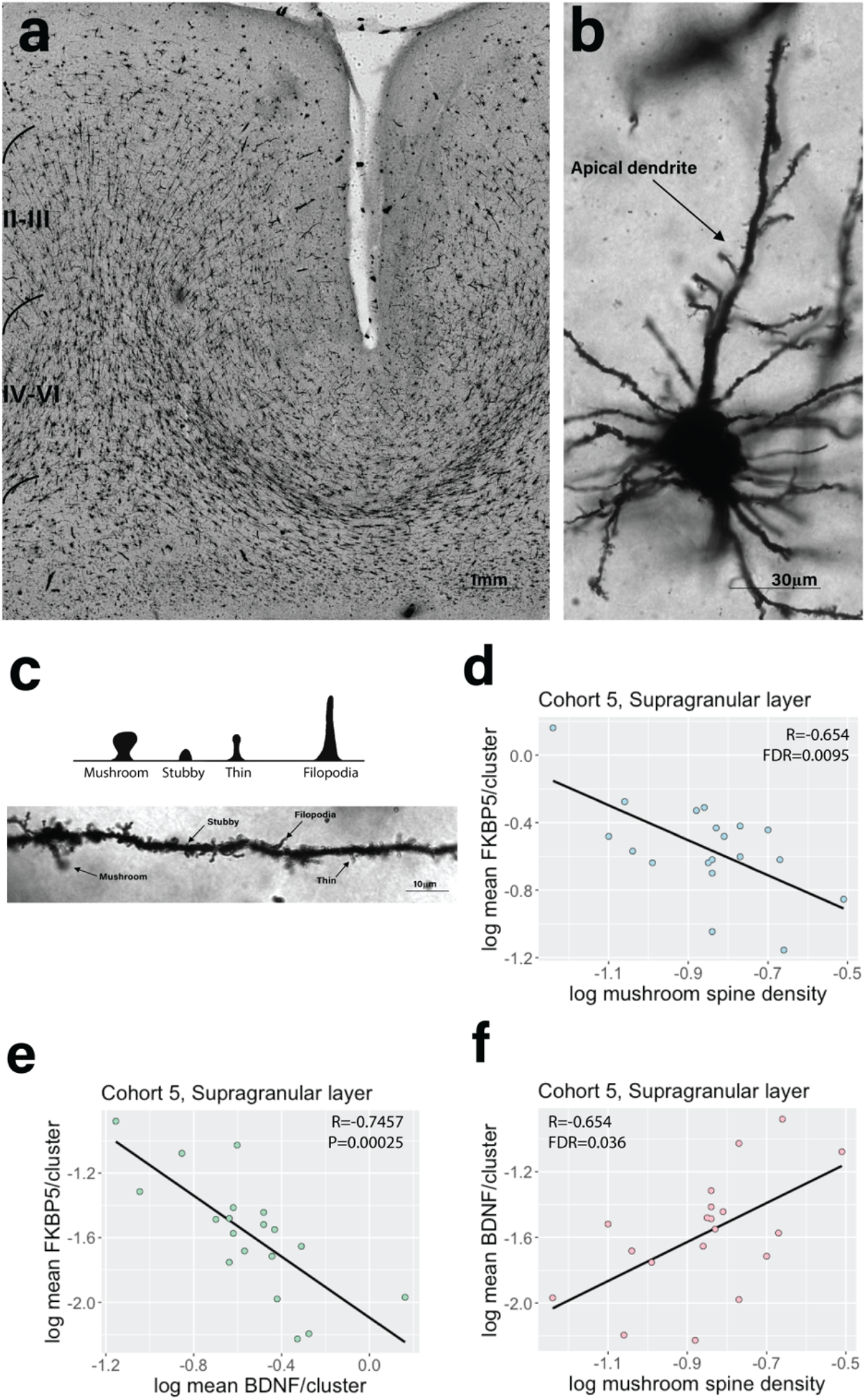
**(a)** Representative image of a cortical section stained with Golgi-Cox. **(b)** Example of an apical dendrite on a pyramidal neuron. **(c)** Classification of different types of dendritic spines: mushroom, stubby, thin and filopodia, both with a diagram (left) and example image of a dendrite (right). Spearman correlations of *FKBP5* gene expression with **(d)** mushroom spine densities and **(e)***BDNF* gene expression, both showing a strong inverse correlation. **(f)** Spearman correlation showing BDNF and mushroom spine density were strongly positively correlated.

Given Brain-Derived Neurotrophic Factor (*BDNF*) is a crucial modulator of dendritic spines [16, 54], and a strong relationship between *FKBP5* and *BDNF* gex levels have been reported [29], we investigated correlations of both *FKBP5* gex with *BDNF* and mushroom spines within the superficial layer neurons in Cohort 5 (*BDNF* UMAPS: Online Resource, Supplementary Figure 4). *BDNF* gex levels were strongly and inversely correlated with *FKBP5* gex levels (R=−0.7457187, *P*=0.00025; Figure 6e), as well as being strongly negatively and specifically correlated with mushroom spine density (R=−0.6541611, *P*=0.002378, FDR=0.036048; Figure 6f). These data provide insight into a possible consequence of elevated *FKBP5* directly in the human brain, indicating a strong relationship between *FKBP5* and mushroom spine density possibly mediated by *BDNF* in supragranular neurons.

## DISCUSSION

With a strong body of evidence from human peripheral blood and preclinical animal studies spurring the development of FKBP51-targeting drugs for psychiatric disorders, it has long been crucial to close the gap on how FKBP51 is cell-type-specifically affected directly in the human brain [31]. In this large-scale postmortem study, we showed that cortical *FKBP5*/FKBP51 mRNA/protein (*FKBP5*/1) expression levels are consistently increased in psychopathology and exacerbated with age, with strikingly higher gene and protein expression levels in individuals with schizophrenia, and depression to a lesser extent. We then traced these effects to the cell-type level to show that they were consistently more pronounced in supragranular excitatory neurons relative to other cell types, and highly associated with cellular morphology. Specifically, we showed that higher levels of FKBP51 inversely correlate with reduced *BDNF* levels and decreased density of mature dendritic spines on these neurons. These findings might have important implications for the dysregulation of cortical circuitry in individuals with schizophrenia and possibly depression, especially those that are older in age. This opens the door for a novel strategy of therapeutic treatment development.

Cognitive deficits are among the most debilitating but difficult to treat symptoms in psychiatric disorders, particularly schizophrenia, and commonly associated with worse prognosis [35]. Deficits in executive functioning, emotional regulation and higher order thinking are attributable to dysfunction of prefrontal cortices, especially the dorsolateral prefrontal cortex, orbitofrontal cortex, and the anterior cingulate cortex. These areas are organised into layers which vary in their cellular composition and connections to other brain areas [9]. The supragranular or superficial layer comprises the upper third of the entire prefrontal cortex thickness, and predominately consists of cortico-cortical connections whereby supragranular excitatory neuron subtypes project to other cortical regions and release glutamate on target neurons, with dendrites being the functional contacts between them [44]. In schizophrenia, differences in the morphology of supragranular neurons are a consistent finding [27]. In particular, reduced dendritic spine densities on supragranular neurons have been previously reported [15, 38, 41]. Our finding that heightened *FKBP5* in these neurons is coupled with lower mushroom spine density suggests there are consequences for circuitry, as the necks of mushroom spines play a crucial role in regulating calcium signalling and the incoming excitatory signal through the synapse, particularly via α-amino-3-hydroxy-5-methyl-4-isoxazolepropionic acid (AMPA) receptors [25]. This is interesting given that in transgenic mice overexpressing the human *FKBP5*, FKBP51 exerts effects on chaperone-mediated recycling of AMPA receptors, with high *FKBP5* accelerating the rate of AMPA recycling and thus altering AMPA receptor trafficking [5]. Mushroom spines are also the most mature form of dendritic spine, and a reduction in this spine type suggests there are more chronic changes in dendritic spine architecture that may impact on long-term potentiation [1]. In support, knock-out of *Fkbp5* in mice has been shown to decrease long-term potentiation and increase presynaptic GABA release [43], and transgenic mice overexpressing human FKBP51 display impaired long-term depression as well as spatial reversal learning and memory [5].

Our finding that *FKBP5* expression is strongly inversely correlated not only with mushroom spine density, but also the crucial spine-modulator *BDNF*, suggests heightened *FKBP5* leads to dysfunctional downstream signalling that may impact synaptic plasticity via *BDNF* [54]. This is consistent with the repeatedly described role of FKBP51 as a molecular hub for many downstream pathways including *BDNF* [58]. For example, we have previously shown that glucocorticoid-induced *FKBP5* expression increases the production of the mature and active form of *BDNF* via epigenetic mechanisms and enhanced secretion of matrix metalloproteinase 9 (MMP9), which was positively coupled to dendritic spine density [11, 29]. The inverse correlation between *FKBP5* and *BDNF* in the current study is likely related to direct effects of FKBP51 on DNMT1, leading to increased methylation of the *BDNF* locus and this decreased expression [11]. Heightened *FKBP5* levels are intertwined with a dysregulated stress response and maladaptive cortisol levels over time [56], with the latter being a common finding in psychopathology [32]. FKBP51 can also be increased via lasting epigenetic disinhibition following exposure to early life adversity, a major common risk factor for psychiatric disease [21, 31]. Elevated glucocorticoid concentrations have been independently shown to cause both increased expression of *FKBP5* [36], as well as atrophy and attrition of dendritic spines [18, 33]. Our group and others have also reported that *FKBP5* levels in the postmortem orbitofrontal cortex are inversely correlated with dendritic mushroom spine density in post-traumatic stress disorder [53] and in mixed psychiatric cases with a history of severely stressful life events [19]. Interestingly, both excitatory and inhibitory supragranular neurons in the dorsolateral prefrontal cortex were recently reported to be vulnerable to the effects of psychopathology in schizophrenia, indicating a core network impairment in this sub-region that may originate from gene × environment interactions during development [2]. Our study may indicate that *FKBP5* is a contributor in this process given *FKBP5* is governed by gene × environment interactions, and its expression is increased via epigenetic mechanisms in the presence of early life adversity [31].

Until now, *FKBP5*/1 cell-type distribution and expression patterns in the human brain have been largely unknown. We show that FKBP51 is highly expressed across virtually all cell-types, consistent with the expression of glucocorticoid receptors which are reportedly ubiquitously expressed [40]. We show that the most pronounced and consistent expression of *FKBP5*/1 is in excitatory neurons in both the supra- and infragranular layers. While *FKBP5* snRNAseq expression appears to be high on microglia and astrocytes, FKBP51 immunostaining in these cells was more inconsistent, with very high staining intensity noted on some cells and undetectable staining on others. Of note, more microglia (and nuclei in total) were captured from cases compared to controls (480 vs 372 respectively [39]), which could contribute to the increased *FKBP5* gex observed overall given microglia appear to be high expressors of *FKBP5* mRNA. It is also possible that microglia-associated *FKBP5* expression may be reflective of increased microglia numbers which is suggested to occur in psychiatric disorders [18]. Regarding neuronal expression, it should be noted that supragranular excitatory neurons are functionally diverse with differing projections in cortical circuitry [37, 55]. Our study did not specifically differentiate between these subtypes. Our immunostaining images suggested that FKBP51 staining intensity on NeuN+ neurons did not vary according to NeuN+ cell morphology, however we acknowledge that 20-35% of supragranular NeuN+ neurons did not express FKBP51. We hypothesise that the lack of expression on certain neurons is more likely to be state-dependent rather than sub-class dependent, given there was no notable sublaminar distribution of FKBP51, and *FKBP5*/1 expression may rapidly fluctuate in response to stress [46].

Previous gene **×** environment studies investigating the effects of the *FKBP5* risk haplotype on the development of psychiatric disorders have reported a cross-diagnostic effect that is not specific to one disorder, but can raise risk to many psychopathologies [31]. Carriers of the *FKBP5* risk haplotype are at an increased risk to develop a psychiatric disorder, but only if exposed to early life adversity [22, 23]. In our study, we found there was no additive effect of the risk haplotype on *FKBP5* expression. However it is important to note that we did not have childhood adversity information for our subjects, although childhood adversity is a common occurrence in individuals with psychiatric disorders [20]. Exposure to childhood adversity is a key component of the genetic and especially epigenetic mechanism that causes disinhibition of *FKBP5* transcription and heightened levels of *FKBP5* [23]. Decreases in DNA methylation in functional *FKBP5* enhancers are observed following exposure to early adversity or exposure to glucocorticoids and accentuated in *FKBP5* risk haplotype carriers [24, 42, 52]. Importantly, also in the human cortex, decreases in DNA methylation were seen in the same enhancers with increasing age [6]. This suggests that enhanced *FKBP5* expression may be differentially driven by different combinations of environmental and genetic risk factors and in relation to age, so it is likely that not all patients are affected by higher *FKBP5* expression. Indeed, our data shows a spread of *FKBP5*/1 expression (Figure 1–2) suggesting that for at least a subset of schizophrenia subjects, particularly those that are older, FKBP51 antagonism may present as an efficacious treatment approach with potential to ameliorate cognitive deficits via effects of dendritic spines as previously shown in cell cultures [29]. So far, data on reliable peripheral or other biomarkers for increased brain *FKBP5* are missing. As larger postmortem cohorts with both information regarding childhood adversity and matching peripheral blood samples are becoming available, this will be important to clarify for clinical translation. As pharmacological targeting of FKBP51 has become more specific [8], another compelling way to study this question could be made possible with the development of positron-emission topography imaging ligands targeting FKBP51, enabling its study in the brain of living people.

This study fills a crucial knowledge gap regarding the cell-type specificity of FKBP51 alterations in the human brain and pinpoints a functional pathway of high relevance to treatment development. Our findings support that FKBP51 antagonists may be most suitable for older individuals with schizophrenia, and possibly depression. Future studies would need to clarify if this is only in a subset of patients that carry the *FKBP5* risk genotype and have been exposed to childhood adversity. This work opens the door for new targeted development of novel drugs that target cortical FKBP51, such as SAFiT2 [29], with the potential to treat cognitive deficits, which are the most debilitating but untreated symptoms of psychiatric disorders, via *BDNF*-mediated synaptic plasticity.

## ACKNOWLEDGEMENTS

We thank the families from the USA, Canada, Germany and Australia who donated brain tissues to this research. We specifically thank the Lieber and Maltz families for their contributions towards the Lieber Institute brain collection. The Douglas-Bell Canada Brain Bank is funded by the Réseau Québécois sur le suicide, le troubles de l’humeur et les troubles associés (FRQS) as well as by Healthy Brains for Healthy Lives (CFREF) and Brain Canada grants to Dr Mechawar and Dr Turecki. The Victorian Brain Bank tissue collection was supported by the National Health and Medical Research Council (NHMRC; Australia; grant number 566967) and the Cooperative Research Centre (CRC) for Mental Health and the Operational Infrastructure Support from the Victorian State Government. We would additionally like to thank Geoff Pavey for his efforts in curating the Australian Victorian Brain Bank and Madhara Udawela for supervising preparation of brain tissues in Cohort 2. Tissues received from the New South Wales Brain Tissue Resource Centre at the University of Sydney were supported by the University of Sydney. Research reported in this publication was supported by the National Institute of Alcohol Abuse and Alcoholism of the National Institutes of Health under Award Number NIAAA012725-15. The content is solely the responsibility of the authors and does not represent the official views of the National Institutes of Health.

## AUTHOR CONTRIBUTIONS

N.Ma. and E.B.B. conceived, designed and managed the work, performed all analyses and drafted the manuscript. J.A., D.C., M.M., A.S.F., T..H, R.T. and A.E.J. were all involved in conceptualising and assisting with the statistical analyses. J.A., D.C., T.H. and C.C. were also involved in substantively drafting and revising the work. S.M., D.K., R.B., K.Z.E., A.R.C., N.C.G., K.H. performed immunoblot and histological analyses, and assisted with interpretation of this data. V.M., N.S.M., K.W., G.R., C.N., M.G., N.G., were involved preparation of omics data for statistical analyses. M.Z., E.S., T.A., P.F., J.E.K., D.R.W., N.Me., A.S., B.D., G.T., and T.M.H. were all involved in collection, dissection and curation of the brain samples used in this study, as well as managing the experiments and sharing the resulting datasets. All authors were involved in the acquisition, analysis and interpretation of the data, and approved the final version of the manuscript.

## DECLARATIONS

Dr Binder is co-inventor on the following patent applications: FKBP5: a novel target for antidepressant therapy. European Patent # EP1687443 B1; Polymorphisms in ABCB1 associated with a lack of clinical response to medicaments. United States Patent # 8030033; Means and methods for diagnosing predisposition for treatment emergent suicidal ideation (TESI). European application number: 08016477.5, international application number: PCT/EP2009/061575. Dr Falkai has been an honorary speaker for AstraZeneca, Bristol Myers Squibb, Eli Lilly, Essex, GE Healthcare, GlaxoSmithKline, Janssen Cilag, Lundbeck, Otsuka, Pfizer, Servier, and Takeda. During the past 5 years, but not presently, Dr Falkai has been a member of the advisory boards of Janssen-Cilag, AstraZeneca, Eli Lilly, and Lundbeck. Dr Schmitt has been an honorary speaker for TAD Pharma and Roche and has been a member of advisory boards for Roche. The remaining authors declare no competing interests or conflicts of interest.

