## Supplementary materials for "Associations of psychiatric disease and aging on *FKBP5* expression converge on cortical supragranular neurons"

Contents

A. **Extended Methods**
   1. Dissection protocols, clinical and demographic details for postmortem cohorts
      a. Supplementary Table 1: Demographics of the LIBD lifetime cohort
      b. Supplementary Table 2: Demographics of the Victorian Brain Bank cohort
      c. Supplementary Table 3: Demographics from the Munich Brain Bank cohort
      d. Supplementary Table 4: Demographics from the Stanley Neuropathology Consortium
      e. Supplementary Table 5: Demographics from the NSW Tissue Resource Centre cohort
      f. Supplementary Table 6: Demographics from the Douglas-Bell Canada Brain Bank cohort
   2. Primer design for real-time quantitative PCR experiments
      a. Supplementary Table 7: RT-qPCR primer design
   3. Antibody validation experiments
      a. Supplementary Table 8: FKBP51 antibodies validated in FKBP51 knock-out cells
      b. Supplementary Figure 1: Validation of FKBP51 antibodies in *FKBP5* knock-out HELA cells using immunoblot
      c. Supplementary Table 9: Summary of antibodies used for immunohistochemistry experiments

B. **Extended Results**
- Supplementary Table 10: Further information about statistical methods used.
- Supplementary Table 11: Sample sizes for genotype analyses
- Supplementary Figure 2: Association of *FKBP5*/1 expression with age in Cohort 2, 3 and 4
- Supplementary Table 12: Schizophrenia-control differences in *FKBP5* gene expression between the major cell-types and excitatory neuron sub-clusters from Cohort 5 using linear regression modelling
- Supplementary Table 13: Depression differences in *FKBP5* gene expression between the major cell-type clusters and excitatory neuron sub-clusters from Cohort 6 using linear regression modelling
- Supplementary Figure 3: Case-control differences in *FKBP5* gene expression in each cell-type cluster from Cohort 5 and Cohort 6
- Supplementary Table 14: Results of Spearman correlations assessing the relationship between *FKBP5* gene expression with age in each cell cluster from Cohort 5
- Supplementary Table 15: Results of Spearman correlations assessing the relationship between *FKBP5* gene expression with age in each cell cluster from Cohort 6
- Supplementary Table 16: Results of Spearman correlations in Cohort 4, assessing the relationship between age and (a) FKBP51 staining intensity on NeuN+ neurons, and (b) the number of FKBP51+ NeuN+ cells in the ACC.
- Supplementary Figure 4. Dimensionality reduction uniform manifold approximation and projection (UMAP) plots depicting (a) *BDNF* expression across cell clusters and (b) bar plot showing average *FKBP5* gene expression per cell-type cluster from Cohort 5.

C. **References**

---

### A. EXTENDED METHODS

#### 1. Dissection protocols, clinical and demographic details for postmortem cohorts

*Postmortem brain Cohort 1*

The LIBD cohort (Cohort 1; Supplementary Table 1) consisted of 252 control subjects ranging from neonate, infant, child, adolescent, and adult age-groups, and 184 schizophrenia (SZ) subjects, 69 bipolar disorder (BPD) and 152 major depressive disorder (MDD) cases. Brains from Cohort 1 were collected at the Clinical Brain Disorders Branch (CBDB), the US National Institute of Mental Health (NIMH), the Northern Virginia and the District of Columbia Medical Examiner's Office (informed consent obtained from legal next of kin for all cases; NIMH Protocol 90-M-0142). Additional fetal, child, and adolescent brain samples were obtained through the National Institute of Child Health and Human Development Brain and Tissue Bank for Developmental Disorders (N01-HD-4-3368 and N01-HD-4_3383). Briefly, DLPFC grey matter from fetal cases was extracted using a dental drill from hemisected 1-1.5cm coronal slabs. For non-fetal cases, Brodmann area 9 and 46 were dissected from the middle frontal gyrus immediately anterior to the genu of the corpus callosum. Dissected tissues were pulverized and stored at -80°C. Detailed information about brain tissue collection and retrieval are available elsewhere (Tao, Davis et al. 2017).

**Supplementary Table 1**. Demographics of the LIBD lifetime cohort

| Descriptive | Group; mean (range)* | | | |
|---|---|---|---|---|
| | CTRL | MDD | BD | SCZ |
| Subjects | 340 | 132 | 59 | 102 |
| Age at death, yr | 30.12 (14 weeks prenatal to 85) | 43.7 (21-64) | 43.7 (21-65) | 46.0 (20-62) |
| Sex, male:female | 228:112 | 79:53 | 32:27 | 69:33 |
| PMI, h | 24.79 (1-90) | 36.4 (5-160) | 29.5 (5-65) | 39.0 (7-142) |
| pH | 5.72 (5.9-7.1) | 6.4 (5.9-7.3) | 6.3 (5.9-6.9) | 6.4 (5.9-7.0) |
| RIN | 8.4 (5-10) | 8.0 (5.1-9.5) | 7.8 (5.0-9.3) | 8.0 (5.4-9.4) |
| Cause of Death†, 1:2:3:4:5 | 2:53:186:0:37 | 2:20:28:79:1 | 2:7:11:39:0 | 3:15:64:20:0 |
| Ethnicity‡ 1:2:3:4 | 145:177:9:9 | 116:11:3:2 | 49:5:2:3 | 55:42:2:3 |

***Abbreviations:*** CTRL, control; MDD, major depressive disorder; BPD, bipolar disorder; SCZ, schizophrenia; PMI, postmortem interval; RIN, RNA integrity number. *Unless otherwise indicated. † 1 = Undetermined, 2 = Accident, 3 = Natural, 4 = Suicide, 5 = Homicide. ‡ 1 = Caucasian, 2 = African American, 3 = Hispanic, 4 = Asian.

*Postmortem brain Cohort 2*

The Victorian brain bank cohort (Cohort 2; Supplementary Table 2) consists of 20 schizophrenia, 20 major depression and 16 bipolar subjects, as well as 20 matched controls. The Diagnostic Instrument for Brain Studies was used to conduct case history reviews to reach a diagnostic consensus using the DSM-IV criteria. Regarding dissection, briefly, BA9 was isolated from the lateral surface of the frontal lobe containing the middle frontal gyrus superior to the inferior frontal sulcus. Tissue blocks were stored at -80°C until processing for downstream applications. The collection and dissection process has been previously described in detail (Scarr, Udawela et al. 2018).

**Supplementary Table 2**: Demographics of the Victorian Brain Bank cohort, postmortem samples of dorsolateral prefrontal cortex (BA9) from the left hemisphere

| Descriptive | Group; mean (range)* | | | |
|---|---|---|---|---|
| | CTRL | MDD | BPD | SCZ |
| Subjects | 20 | 20 | 16 | 20 |
| Age at death (years) | 59.2 (32-80) | 85.0 (27-87) | 59.4 (31-79) | 56.4 (30-82) |
| Sex (male:female) | 11:9 | 11:9 | 8:8 | 11:9 |
| Suicide (no:yes) | 19:1 | 3:17 | 10:6 | 14:6 |
| Brain pH | 6.3 (5.9-6.7) | 6.5 (5.6-6.9) | 6.3 (6.0-6.5) | 6.3 (5.5-6.6) |
| PMI (hours) | 43.0 (17-72) | 43.4 (11-72) | 38.7 (8-63) | 44.2 (20-66) |
| Brain weight (grams) | 1354 (1140-1655) | 1285 (1030-1573) | 1244 (952-1630) | 1337 (1110-1659) |

***Abbreviations:*** CTRL, control; MDD, major depressive disorder; BPD, bipolar disorder; SCZ, schizophrenia; PMI, postmortem interval; RIN, RNA integrity number. *Unless otherwise indicated

*Postmortem brain Cohort 3*

The Munich Neurobiobank cohort (Cohort 3; Supplementary Table 1b) consists of 24 control subjects with ages evenly distributed over adulthood, from 37 to 88 years of age. Clinical records were provided by relatives and general practitioners and all assessments and postmortem evaluations were conducted in accordance with the Ethics Committee of the Faculty of Medicine, University of Heidelberg, Germany. Controls had no history of alcohol or drug abuse, severe physical or psychiatric illness. Tissue blocks (~1-1.5cm$^2$) were dissected from the anterior-most point of the superior frontal gyrus, corresponding to BA9. Braak neuropathological examinations were staged at ≤2 for all subjects. Tissue blocks were stored at -80°C until processing for downstream applications.

**Supplementary Table 3**: Demographics from Munich Brain Bank, postmortem control cohort with samples from the dorsolateral prefrontal cortex (BA9)

| Descriptive | Group; mean (range)* |
|---|---|
| | **CTRL** |
| Subjects | 24 |
| Age at death (years) | 65.5 (37-88) |
| Sex (male:female) | 13:11 |
| Hemisphere (left:right) | 11:13 |
| PMI (hours) | 32.6 (13-71) |
| RIN | 6.1 (3.0-8.3) |

***Abbreviations:*** CTRL, control; MDD, major depressive disorder; BPD, bipolar disorder; SCZ, schizophrenia; PMI, postmortem interval; RIN, RNA integrity number. *Unless otherwise indicated

*Postmortem brain Cohort 4*

The Stanley Neuropathology Consortium (Cohort 4, Supplementary Table 4) consists of 60 adult subjects (schizophrenia, major depression, bipolar disorder and matched controls, n=15/group). Samples were matched for age, sex, race, and PMI. Detailed clinical and demographic information regarding these cohorts have been published previously (Torrey, Webster et al. 2000).

**Supplementary Table 4**: Demographics from the Stanley Neuropathology consortium, postmortem control cohort with samples from the anterior cingulate cortex (BA24)

| Descriptive | Group; mean (range)* | | | |
|---|---|---|---|---|
| | **CTRL** | **MDD** | **BPD** | **SCZ** |
| Subjects | 15 | 15 | 15 | 15 |
| Age at death (years) | 48.1 (29-68) | 46.5 (30-65) | 42.3 (25-61) | 44.2 (25-62) |
| Age of onset | - | 33.9 (11-54) | 21.5 (7-39) | 23.2 (13-42) |
| Sex (male:female) | 9M, 6F | 9M, 6F | 9M, 6F | 9M, 6F |
| PMI (hours) | 23.7 (8-42) | 27.5 (7-47) | 32.5 (13-62) | 33.7 (12-61) |
| FST (days) | 338.27 (31-774) | 434 (86-931) | 620 (224-836) | 621.13 (68-938) |
| pH | 6.3 (5.8-6.6) | 6.2 (5.6-6.5) | 6.2 (5.8-6.5) | 6.1 (5.8-6.6) |

***Abbreviations:*** CTRL, control; MDD, major depressive disorder; BPD, bipolar disorder; FST, freezer storage time; SCZ, schizophrenia; PMI, postmortem interval. *Unless otherwise indicated

*Postmortem brain Cohort 5*

The NSW Brain Tissue Resource Centre cohort (Cohort 5, Supplementary Table 5) consists of a total of 36 cases with schizophrenia, and 33 matched controls, and tissues were derived from the orbitofrontal cortex, BA11. 45 individuals were male and 24 female, and samples matched by age, PMI and RIN. Postmortem brain tissue was collected at the New South Wales Brain Tissue Resource Centre at the University of Sydney which is supported by the University of Sydney. Research reported in this publication was supported by the National Institute of Alcohol Abuse and Alcoholism of the National Institutes of Health under Award Number NIAAA012725-15. Informed consent was given by all donors or their next of kin for brain autopsy. Groups were matched according to psychiatric diagnoses, postmortem interval (PMI), age at death, and RNA integrity number (RIN) (Supplementary Table 5).

**Supplementary Table 5**. Demographics from NSW Brain Tissue Resource Centre postmortem cohort with samples from the orbitofrontal cortex (BA11).

| | Group (mean±standard deviation*) | |
|---|---|---|
| **Descriptive** | **Control** | **SZ** |
| Subjects | 33 | 36 |
| Age at death (years) | 57±4.32 | 41.69±4.76 |
| Sex (male:female) | 17:0 | 17:0 |
| Cause of death | Natural death (11), accident (6) | Suicide (17) |
| Brain pH | 6.49±0.06 | 6.60±0.07 |
| PMI (hours) | 34.01±4.94 | 41.69±4.76 |
| RIN | 6.16 | 6.47 |

***Abbreviations:*** PMI, postmortem interval; SZ, schizophrenia; RIN, RNA integrity number. *Unless otherwise indicated

*Postmortem brain Cohort 6*

The Douglas-Bell Canada Brain Bank cohort (Cohort 6, Supplementary Table 6) consists of a total of 17 cases with major depressive disorder who died by suicide, and 17 matched controls. All cases were males, and samples matched by age, PMI and RIN. Cause of death was determined by the Quebec Coroner's office, and frozen grey matter from BA9 isolated by trained neuroanatomists. Psychological autopsies were performed using proxy-based interviews. Cases met criteria for MDD and died by suicide, and controls were individuals who died suddenly and did not have evidence of any axis I disorders. The study was approved by the Douglas Hospital Research Ethics Board. Written informed consent was received from the next of kin for each individual.

**Supplementary Table 6**. Demographics from Douglas-Bell Canada Brain Bank postmortem cohort with samples from the dorsolateral prefrontal cortex (BA9).

| | Group (mean±standard deviation*) | |
|---|---|---|
| **Descriptive** | **Control** | **MDD** |
| Subjects | 17 | 17 |
| Age at death (years) | 38±4.32 | 41.69±4.76 |
| Sex (male:female) | 17:0 | 17:0 |
| Cause of death | Natural death (11), accident (6) | Suicide (17) |
| Brain pH | 6.49±0.06 | 6.60±0.07 |
| PMI (hours) | 34.01±4.94 | 41.69±4.76 |
| RIN | 6.16 | 6.47 |

***Abbreviations:*** MDD, major depressive disorder; PMI, postmortem interval; RIN, RNA integrity number. *Unless otherwise indicated

### 2. Primer design for real-time quantitative PCR experiments

**Supplementary Table 7.** RT-qPCR primer design

| | Details | | FKBP5 target | Region |
|---|---|---|---|---|
| 1 | IDT Ref | Hs.PT.58.813038 (all transcripts Exon 11-12) | All transcripts | Exon 11-12 |
| | Probe | /56-FAM/CTG TTG AAT /ZEN/GCT GTG ACA AGG CCC /3IABkFQ/ | | |
| | Primer 1 | ATG TGC TAC CTG AAG CTT AGA G | | |
| | Primer 2 | CCC TCC TAT ACA AGC CTT TCT C | | |
| 2 | IDT Ref | Hs.PT.58.20523859 (all transcripts Exon 5-6) | All transcripts | Exon 5-6 |
| | Probe | /56-FAM/AGAGATATG/ZEN/CCATTTACTGTGCAAACCAGA/3IABkFQ/ | | |
| | Primer 1 | GAACCATTTGTCTTTAGTCTTGGC | | |
| | Primer 2 | CGAGGGAATTTTAGGGAGACTG | | |
| 3 | IDT Ref | Hs.PT.39a.22214836 (GAPDH (NM_002046) Exon 2-3) | GAPDH | Exon 2-3 |
| | Probe | /56-FAM/AAGGTCGGA/ZEN/GTCAACGGATTTGGTC/3IABkFQ/ | | |
| | Primer 1 | ACATCGCTCAGACACCATG | | |
| | Primer 2 | TGTAGTTGAGGTCAATGAAGGG | | |
| 4 | IDT Ref | Hs.PT.39a.22214847 (ACTB Exon 1-2) | ACTB | Exon 1-2 |
| | Probe | | | |
| | Primer 1 | | | |
| | Primer 2 | | | |

#### 3. Antibody validation

To validate the FKBP51 antibodies used in our study, immunoblots were performed in FKBP51 knock out (KO) cells. The KO cells were generated from the SH-SY5Y human neuroblastoma cell line using CRISPR-Cas9 (Martinelli et al.). Considering FKBP51 is lowly expressed at baseline, the synthetic glucocorticoid receptor agonist dexamethasone was also added to induced FKBP5 expression and increase visible expression of FKBP51 protein (Figure S1). Protein was extracted from cell lysates and 20ug was loaded for immunoblot analyses. FKBP51 antibodies (Supplementary Table 5) were applied (on individually run membranes) at 1:1000, and membranes were imaged according to the general methods.

**Supplementary Table 8.** FKBP51 antibodies validated in FKBP51 knock-out cells

| Name | Type | Concentration | Number | Company |
|---|---|---|---|---|
| Rabbit anti-FKBP51 | polyclonal | 1:1000 | #8245S | Cell Signaling, Danvers, MA, USA |
| Rabbit anti-FKBP51 | polyclonal | 1:1000 | #A301-429A | Bethyl Laboratories, Montgomery, TX, USA |
| Mouse anti-FKBP51 | polyclonal | 1:1000 | #D-4 | Santa Cruz Biotechnology, Dallas, TX, USA |

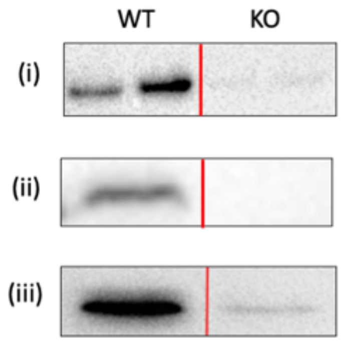

**Supplementary Figure 1.** Validation of FKBP51 antibodies in *FKBP5* knock-out HELA cells using immunoblot. All FKBP51 antibodies ([i] Cell Signalling #8245S, [ii] Bethyl #A301-429A, [iii] Santa Cruz #D-4) showed specific expression in wildtype (WT) cells but not *FKBP5* knock-out (KO) cells, indicating their specificity.

**Supplementary Table 9**. Summary of antibodies used for immunohistochemistry experiments

| | Antibody | Concentration | Catalog Number | Supplier; Country |
|---|---|---|---|---|
| Primary Abs | Rabbit anti-FKBP5 | 1:200 | #8245S | Cell Signalling Technologies; MA, USA |
| | Goat anti-GAD1 | 1:26 | #AF2086 | R&D Systems; MN, USA |
| | Mouse anti-NeuN | 1:100 | #MAB377 | Sigma-Aldrich; MA, USA |
| | Mouse anti-FKBP5 | 1:200 | #sc-271547 | Santa Cruz; TX, USA |
| | Rabbit anti-TMEM119 | 1:100 | #ab185333 | Abcam, Cambridge UK |
| | Chicken anti-GFAP | 1:500 | #AB5541 | Sigma-Aldrich; MA, USA |
| Secondary Abs | Donkey anti-rabbit 488 | 1:500 | #A32790 | Invitrogen; MA, USA |
| | Donkey anti-goat 594 | 1:500 | #A32758 | Invitrogen; MA, USA |
| | Donkey anti-mouse 647 | 1:500 | #A32787 | Invitrogen; MA, USA |
| | Donkey anti-chicken 647 | 1:300 | #703-605-155 | Jackson Immuno Research; PA, USA |

### B. EXTENDED RESULTS

**Supplementary Table 10.** Summary of the research questions and statistical methods applied in this study.

| Research question | Statistical method | Statistical model | Covariates (only selected if significant in the model) | Cohort and sample size | Experimental Method | Result |
|---|---|---|---|---|---|---|
| Does *FKBP5* gene expression significantly differ between cases (grouped and independent diagnosis) and controls? | Linear model | lm(gex~cov+diagnosis) | Age, sex, PMI, pH, qSVA†, cell type* | Cohort 1 (14 years+ only)<br>n=180 controls<br>n=121 schizophrenia<br>n=144 major depression<br>n=63 bipolar | Bulk RNA sequencing | *FKBP5* expression levels were significantly heightened in cases, driven by schizophrenia subjects. |
| | | | Age, sex, PMI, pH, suicide, RIN, type of death, experimental batch variables | Cohort 2<br>n=62 controls<br>n=68 schizophrenia<br>n=24 major depression<br>n=15 bipolar | Microarray | |
| Does FKBP51 protein expression significantly differ between cases (grouped and independent diagnosis) and controls? | Linear model | lm(gex~cov+diagnosis) | Age, sex, PMI, pH, suicide, RIN, type of death | Cohort 2<br>n=20 controls<br>n=20 schizophrenia<br>n=20 major depression<br>n=16 bipolar disorder | Western blot | FKBP51 expression levels were significantly heightened in cases, driven by schizophrenia subjects. |
| | | | Age, sex, PMI, pH | Cohort 4<br>n=15 controls<br>n=15 schizophrenia<br>n=15 major depression<br>n=15 bipolar disorder | | |
| Do *FKBP5* gene expression and FKBP51 protein expression correlate? | Spearman's correlation | cor(gex,protein expression) | - | Cohort 2<br>n=20 controls<br>n=20 schizophrenia<br>n=20 major depression<br>n=16 bipolar disorder | Bulk RNA sequencing and western blot | There is a a strong, positive correlation between *FKBP5* gex and FKBP51 protein levels across all subjects. |
| Is there an additive effect of the rs1360780 risk genotype on case status causing further increased *FKBP5* gene expression? | Linear model | lm(gex~cov+SNP+diagnosis) | Age, sex, PMI, pH, qSVA†, cell type* | Cohort 1 (14 years+ only)<br>n=108/64 T/C controls<br>n=69/45 T/C schizophrenia<br>n=78/62 T/C major depression<br>n=32/28 T/C bipolar | Bulk RNA sequencing and SNP genotyping | No additive effect of SNP risk genotype is present. |
| Are *FKBP5* gene expression levels correlated with age? | Spearman's correlation | cor(gex, age) | - | Cohort 1 (14 years+ only)<br>n=180 controls | Bulk RNA sequencing | Strong correlation of *FKBP5* gene expression with age in neurotypical controls. |
| | | | | Cohort 2<br>n=62 controls | Microarray | |
| | | | | Cohort 3<br>n=24 controls | qPCR (two probes) | |

| | | | | | | |
|---|---|---|---|---|---|---|
| Are FKBP51 protein expression levels correlated with age? | Spearman's correlation | cor(protein expression, age) | - | Cohort 2<br>n=20 controls<br>Cohort 3<br>n=24 controls<br>Cohort 4<br>n=15 controls | Western blot | Strong correlation of FKBP51 protein with age in neurotypical controls. |
| Are the effects of aging on *FKBP5* gene expression further increased in psychiatric disorders? | sm.ancova | sm.ancova(gex~cov+diagnosis) | Age, sex, race, RIN, PMI, qSVA, cell type*<br>Age, sex, PMI, pH, suicide, RIN, type of death, experimental batch variables | Cohort 1 (14 years+ only)<br>n=180 controls<br>n=121 schizophrenia<br>n=144 major depression<br>n=63 bipolar<br>Cohort 2<br>n=62 controls<br>n=68 schizophrenia<br>n=24 major depression<br>n=15 bipolar | Bulk RNA sequencing<br>Microarray | In Cohort 1, the *FKBP5* ageing trajectory was significantly heightened in schizophrenia subjects compared to controls.<br><br>In Cohort 2, a similar trend was seen but it did not reach statistical significance (*P*=0.056) |
| Do the case-control differences in *FKBP5* gene expression vary according to cell-type? | Linear model | lm(gex~cov+diagnosis) | Age, PMI, pH, RIN, library prep batch, sequencing batch<br>Age, PMI, pH, RIN, sequencing batch | Cohort 5<br>n=33 controls<br>n=36 schizophrenia<br>Cohort 6<br>n=17 controls<br>n=17 depression | Single-nucleus RNA sequencing<br><br>Comparison in each major cell-type cluster and excitatory neuron subcluster based on cortical layer location. | In schizophrenia, *FKBP5* gene expression was increased in all cell types in the major cell-type clusters except interneurons, with strong effects seen in excitatory cell types. In the excitatory neuron cortical-layer subclusters, *FKBP5* gene expression was higher only in the supragranular layer.<br><br>In depression, the only case-control difference in *FKBP5* geneexpression was found in the excitatory neuron group and supragranular neurons specifically but neither survived correction for multiple cell-type comparisons. |
| Is there an increase in FKBP51 protein expression on supragranular excitatory neurons in schizophrenia? | Linear model | lm(protein expression~cov+diagnosis) | Age, sex, PMI, brain weight, pH, freezer storage time, subject | Cohort 4<br>n=15 controls<br>n=15 schizophrenia<br>n=15 major depression<br>n=15 bipolar disorder<br><br>To improve power, analyses were performed on count and single-cell staining intensity data for >12,000 NeuN+ neurons. | Immunohistochemistry | In schizophrenia, FKBP51 staining intensity was higher in the supgranular layer of the cortex but not the deep layer. In depression, FKBP51 staining intensity was lower in the supgranular layer of the cortex but not the deep layer. There was no differences seen in bipolar disorder. |
| Is elevated superficial neuron *FKBP5* gene expression levels in schizophrenia correlated with dendritic spine architecture (mushroom, stubby, filopodia and thin spines)? | Spearman's correlation | cor(gex, spine density)<br>cor(gex 1, gex 2) | - | Cohort 5<br>n=8 control<br>n=11 schizophrenia | Single-nucleus RNA sequencing<br><br>Golgi-Cox staining | *FKBP5* gene expression levels were specifically, highly and negatively correlated with mushroom spine density.<br><br>*BDNF* gene expression levels were strongly and inversely correlated with *FKBP5* gene expression levels and strongly and specifically correlated with mushroom spines. |

| | | | | | | |
|---|---|---|---|---|---|---|
| Are *FKBP5*/1 mRNA/protein expression levels correlated with age specifically in supragranular neurons? | Spearman's correlation | cor(gex, age)<br>cor(protein expression, age) | - | Cohort 5 (cases and controls combined)<br>n=33 controls<br>n=36 schizophrenia | Single-nucleus RNA sequencing | Age was correlated with *FKBP5* gene expression in all the major cell-type clusters. The most striking correlation was in the excitatory neuron cluster. In the excitatory neuron subclusters, the strongest aging effect was in the supragranular excitatory neurons. |
| | | | | Cohort 6 (cases and controls combined)<br>n=17 controls<br>n=17 depression | Single-nucleus RNA sequencing | Age was strongly correlated with *FKBP5* gene expression in the excitatory neuron cluster. The aging effect was strongest in, and specific to, cortical layer 2-4 excitatory neurons. |
| | | | | Cohort 3<br>n=24 controls | RNAscope | Positive correlation of age and *FKBP5* transcript dots in the superficial layers but not the deep layers. |
| | | | | Cohort 3<br>n=24 controls | Immunohistochemistry | Positive correlation of age on FKBP51 protein expression in NeuN+ neurons was also specific to the superficial but not deep cortical layers. |
| Are the age effects on *FKBP5*/1 mRNA/protein expression levels in supragranular neurons pronounced in cases? | Spearman's correlation | cor(protein expression, age) | - | Cohort 4<br>n=15 controls<br>n=15 schizophrenia<br>n=15 major depression<br>n=15 bipolar disorder<br><br>To improve power, analyses were performed on count and single-cell staining intensity data for >12,000 NeuN+ neurons. | Immunohistochemistry | Schizophrenia subjects showed the strongest correlation of age on the number of FKBP51+ neurons specific to the superficial but not deep cortical layers. MDD and BP subjects showed a weak positive correlation of age and number of FKBP51+ cells only in the deep layers.<br><br>Schizophrenia subjects showed the strongest positive correlation of FKBP51+ neuronal staining intensity with age, in both the superficial and deep layers.<br><br>In MDD, FKBP51+ neuronal staining intensity was positively correlated with age only in the deep layers. In BP, the effects of age were present in both the superficial and deep layers.<br><br>In controls, there was no correlation of age on the numbers of FKBP51+ neurons or neuronal staining intensity in neither the deep nor superficial layers |

* To account for potential confounding of different cell types, we used cell type variables derived from the RNA deconvolution model previously described (Darmanis, Sloan et al. 2015)
† To remove residual confounding by RNA degradation, we used the quality surrogate variable analysis (qSVA) framework previously described (Jaffe, Tao et al. 2017)

**Supplementary Table 11.** Sample sizes for genotype analyses

| **Rs1360780,** Sample size (n) | | |
|---|---|---|
| | T carrier | CC |
| Control | 108 | 64 |
| Schizophrenia | 69 | 45 |
| Major Depression | 78 | 62 |
| Bipolar Disorder | 32 | 28 |

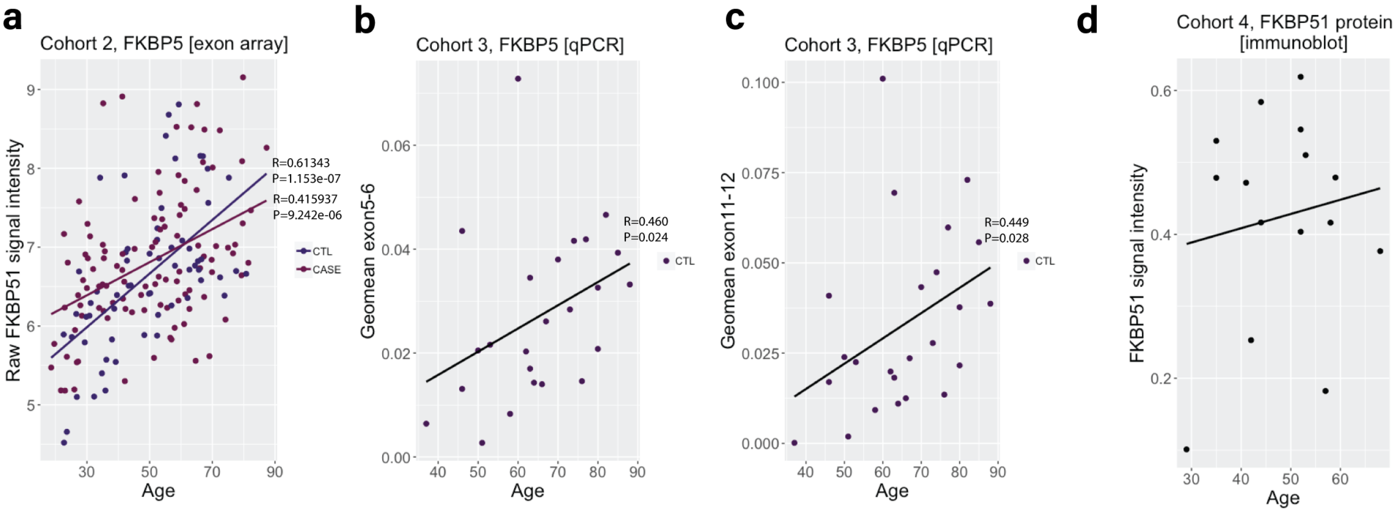

**Supplementary Figure 2. Association of *FKBP5/*1 expression with age in Cohort 2, 3 and 4. (a)** Association of *FKBP5* gene expression with age in Cohort 2. *FKBP5* was positively and significantly associated with age in both cases and controls (CTL), with heightened expression at older ages in cases vs controls. The difference in case vs control *FKBP5* aging trajectory, measured using sm.ancova, did not reach statistical significance ($P$=0.056). **(b-c)** Results of quantitative PCR experiments assessing the correlation between *FKBP5* expression with age in Cohort 3. *FKBP5* was significantly associated with age with two independent probes targeting all transcripts either at (**b**) *FKBP5* exon 5-6 or (**c**)11-12. Cohort refers to the cohorts used in each analysis, as detailed in Table 1. **(d)** Association of FKBP51 protein expression with age in Cohort 4. *FKBP5* was not correlated with age, although it is important to note that there were few subjects over 60 in this sample (n=1).

**Supplementary Table 12.** Schizophrenia vs control differences in *FKBP5* gene expression between the major cell clusters and excitatory neuron sub-clusters from Cohort 5, using linear regression modelling. Results of the coefficients for diagnoses derived from the linear regressions (*lm* function) are presented. Nominal and FDR-corrected P values for multiple comparisons across cell types are reported.

| **COHORT 5**<br>**Major/sub-cluster name** | **Cortical layer(s)** | **t-value** | $P_{nom}$ | $P_{FDR}$ |
|---|---|---|---|---|
| Oligodendrocytes | - | 3.128 | 0.002701 | **0.006302333** |
| Astrocytes | - | 3.135 | 0.002644 | **0.006302333** |
| Oligodendrocyte progenitor cell | - | 3.452 | 0.00102 | **0.006302333** |
| Inhibitory neurons | - | 1.937 | 0.05740 | 0.057400000 |
| Endothelial cells | - | 2.745 | 0.00794 | **0.011116000** |
| Microglia | - | 2.578 | 0.012354 | **0.014413000** |
| Excitatory | - | 2.969 | 0.00426 | **0.007455000** |
| *Posthoc:* | | | | |
| Ex 2 | 2-3 | **3.256** | **0.00186** | **0.00930000** |
| Ex 3 | 3-5 | 2.123 | 0.03793 | 0.06321667 |
| Ex 4 | 4-6 | 2.265 | 0.02721 | 0.06321667 |
| Ex 5 | 4-6 | 1.956 | 0.0551 | 0.06887500 |
| Ex 6 | 5-6 | 1.410 | 0.163877 | 0.16387700 |

**Supplementary Table 13.** Depression vs control differences in *FKBP5* gene expression between excitatory neuron sub-clusters from Cohort 6 using linear regression modelling. Results of the coefficients for diagnoses derived from the linear regressions (*lm* function) are presented. Nominal and FDR-corrected P values for multiple comparisons across cell types are reported.

| **COHORT 6**<br>**Major/sub-cluster name** | **Cortical layer(s)** | **t-value** | $P_{nom}$ | $P_{FDR}$ |
|---|---|---|---|---|
| Astrocytes | - | 1.079 | 0.29058 | 0.5749333 |
| Endothelium | - | 0.328 | 0.744114 | 0.744114 |
| Microglia | - | 1.335 | 0.18717 | 0.5749333 |
| Oligodendrocytes | - | -0.731 | 0.471 | 0.5749333 |
| Oligodendrocyte progenitor cell | - | 0.766 | 0.4445 | 0.5749333 |
| Inhibitory | - | 0.695 | 0.4928 | 0.5749333 |
| Excitatory | - | **-1.776** | **0.0267** | 0.1869 |
| *Posthoc:* | | | | |
| Ex 2 | 5 | 0.066 | 0.948 | 0.9480000 |
| Ex 3 | 4-5 | 0.478 | 0.638 | 0.7443333 |
| Ex 4 | 6 | -0.687 | 0.4990 | 0.6986000 |
| Ex 6 | 4-6 | 0.729 | 0.4723 | 0.6986000 |
| Ex 7 | 4-6 | -0.978 | 0.33673 | 0.6986000 |
| Ex 8 | 5-6 | -1.596 | 0.1226 | 0.4291000 |
| Ex 10 | 2-4 | **-2.190** | **0.0364** | 0.2548000 |

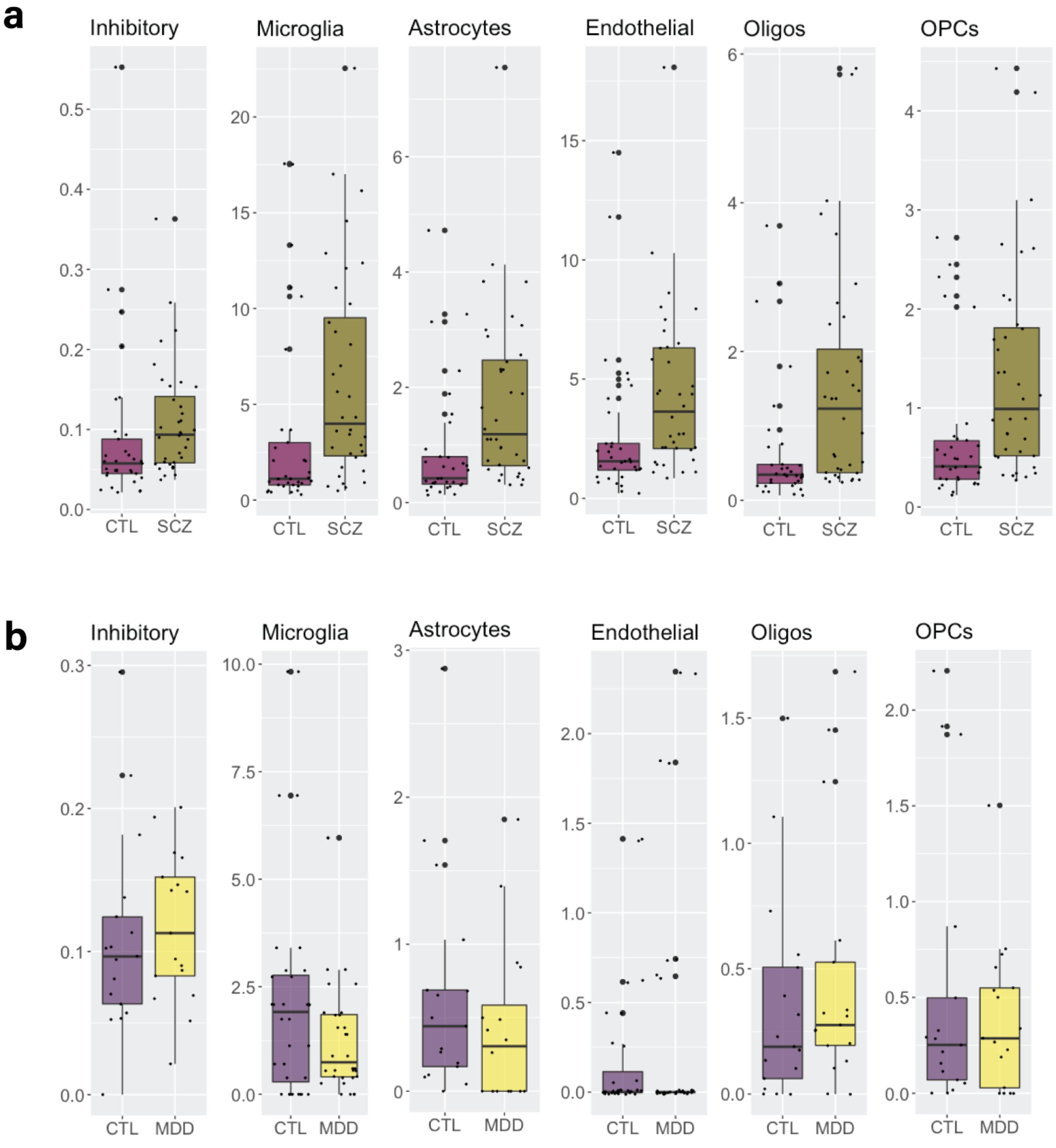

**Supplementary Figure 3.** Boxplots showing snRNAseq *FKBP5* gene expression in each individual major cell type cluster (labelled on the top of each plot) from (**a**) Cohort 5 and (**b**) Cohort 6. In schizophrenia, there was a significant difference (P<0.05) in all groups except the inhibitory interneurons (see Supplementary Table 12). In depression, there was no significant difference in *FKBP5* mRNA expression in depression in any of these cell-types. Statistical output is provided in Supplementary Tables 11. Y axis represents mean *FKBP5* gene expression per cluster. *Abbreviations:* CTL, control; Inhibitory, inhibitory neurons, MDD, major depression; oligos, oligodendrocytes; OPCs, oligodendrocyte progenitor cells.

**Supplementary Table 14**. Results of Spearman correlations assessing the relationship between *FKBP5* gene expression with age in each cell cluster from Cohort 5 (schizophrenia and controls combined). Nominal and FDR-corrected P values are reported.

| **Major/sub-cluster name** | **Cortical layer(s)** | **R-value** | **$P_{nom}$** | **$P_{FDR}$** |
|---|---|---|---|---|
| Oligodendrocytes | - | **0.3503878** | **0.003162** | **0.0037180000** |
| Astrocytes | - | **0.3447741** | **0.003718** | **0.0037180000** |
| Oligodendrocyte progenitor cell | - | 0.3747076 | 0.001513 | **0.0035303333** |
| Inhibitory neurons | - | 0.4046593 | 0.0005633 | **0.0019715500** |
| Endothelial cells | - | 0.3533501 | 0.002899 | **0.0037180000** |
| Microglia | - | 0.3478827 | 0.0034 | **0.0037180000** |
| Excitatory neurons | - | **0.5537049** | **8.016e-07** | **0.0000056112** |
| *Posthoc:* | | | | |
| **Ex 2** | **2-3** | **0.555395** | **7.3e-07** | **0.00000365** |
| **Ex 3** | **3-5** | **0.2683298** | **0.0258** | **0.02580000** |
| **Ex 4** | **4-6** | **0.2930017** | **0.01455** | **0.01818750** |
| **Ex 5** | **4-6** | **0.4972975** | **1.38e-05** | **0.00003450** |
| **Ex 6** | **5-6** | **0.3013647** | **0.01186** | **0.01818750** |

**Supplementary Table 15**. Results of Spearman correlations assessing the relationship between *FKBP5* gene expression with age in each cell cluster from Cohort 6 (depression and controls combined). Nominal and FDR-corrected P values are reported.

| **Major/sub-cluster name** | **Cortical layer(s)** | **R-value** | **$P_{nom}$** | **$P_{FDR}$** |
|---|---|---|---|---|
| Oligodendrocytes | - | **0.4011025** | **0.0187** | **0.0479** |
| Astrocytes | - | **0.4015578** | **0.0205** | **0.0479** |
| Oligodendrocyte progenitor cell | - | 0.3309074 | 0.0559 | 0.0979 |
| Inhibitory neurons | - | 0.2962397 | 0.0889 | 0.1245 |
| Endothelial cells | - | 0.1257452 | 0.4786 | 0.5584 |
| Microglia | - | 0.09823437 | 0.5927 | 0.5927 |
| Excitatory neurons | - | **0.6638643** | **1.87E-05** | **1.31E-04** |
| *Posthoc:* | | | | |
| Ex 2 | 5 | 0.1197 | 0.5138 | 0.7193 |
| Ex 3 | 4-5 | 0.3377615 | 0.0849 | 0.1827 |
| Ex 4 | 6 | 0.3023 | 0.1044 | 0.1827 |
| Ex 6 | 4-6 | 0.3122614 | 0.0722 | 0.1827 |
| Ex 7 | 4-6 | 0.06969655 | 0.6953 | 0.7267 |
| Ex 8 | 5-6 | 0.06322 | 0.7267 | 0.7267 |
| **Ex 10** | **2-4** | **0.69978** | **4.11E-06** | **2.88E-05** |

**Supplementary Table 16**. Results of Spearman correlations in Cohort 4, assessing the relationship between age and (a) FKBP51 staining intensity on NeuN+ neurons, and (b) the number of FKBP51+ NeuN+ neurons in the ACC.

| | **Supragranular neurons** | | **Infragranular neurons** | |
|---|---|---|---|---|
| **(a)** Age vs FKBP51 staining intensity on NeuN+ neurons | | | | |
| | R value | P (nominal) value | R value | P (nominal) value |
| All subjects | -0.143087 | 0.2973 | -0.1163699 | 0.3975 |
| Controls | -0.01991194 | 0.9461 | -0.1902697 | 0.5147 |
| Schizophrenia | -0.04410167 | 0.881 | -0.01764067 | 0.9523 |
| Major Depression | -0.2970299 | 0.3024 | -0.2816283 | 0.3293 |
| Bipolar Disorder | -0.4536761 | 0.1194 | 0.008298954 | 0.9785 |
| (b) Age vs the number of FKBP51+ NeuN+ neurons | | | | |
| | R value | P (nominal) value | R value | P (nominal) value |
| All subjects | **0.2735827** | **0.04327** | 0.06430252 | 0.6409 |
| Controls | 0.4589846 | 0.09876 | 0.2893598 | 0.3157 |
| Schizophrenia | **0.6270845** | **0.01638** | -0.08425741 | 0.7746 |
| Major Depression | -0.1283198 | 0.662 | -0.03311266 | 0.9105 |
| Bipolar Disorder | 0.2402787 | 0.4291 | 0.1448468 | 0.6368 |

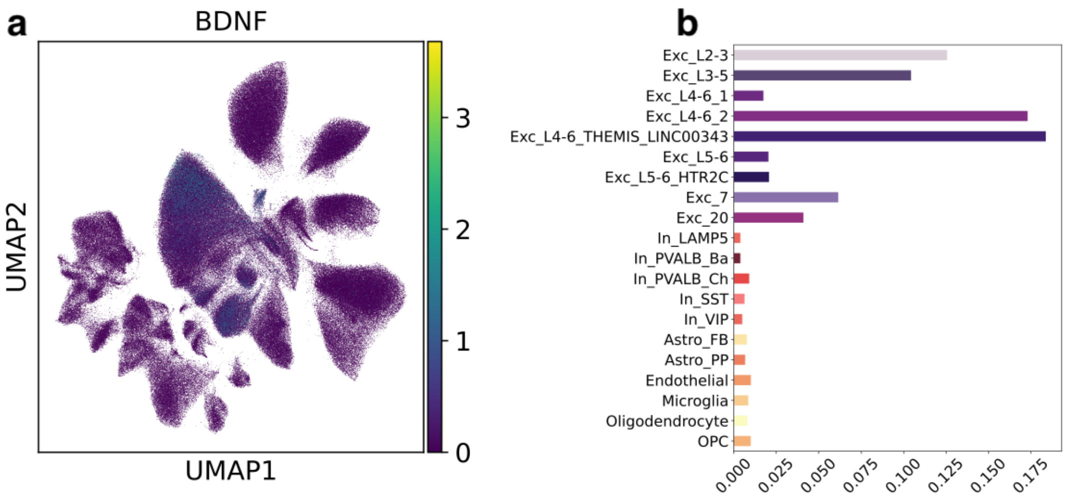

**Supplementary Figure 4.** Dimensionality reduction uniform manifold approximation and projection (UMAP) plots depicting (a) *BDNF* expression across cell clusters and (b) bar plot showing average *FKBP5* gene expression per cell-type cluster from Cohort 5.
